# Single-molecule localization microscopy reveals the ultrastructural root constitution of distal appendages in expanded mammalian centrioles

**DOI:** 10.1101/2022.07.21.500782

**Authors:** Ting-Jui Ben Chang, Jimmy Ching-Cheng Hsu, T. Tony Yang

## Abstract

Distal appendages (DAPs) are vital in cilia formation, mediating vesicular and ciliary docking to the plasma membrane during early ciliogenesis. Although numerous DAP proteins arranging a nine-fold symmetry have been studied using superresolution microscopy analyses, the extensive ultrastructural understanding of the DAP root structure developing from the centriole wall remains elusive owing to insufficient resolution. Here, we proposed a pragmatic imaging strategy for two-color single-molecule localization microscopy of swellable mammalian DAP proteins. Importantly, our imaging workflow enables us to push the resolution limit of a light microscope well close to an electron microscopy level, thus achieving an unprecedented λ/200 mapping precision inside intact cells. Upon this workflow, we unravel the ultra-resolved higher-order protein complexes of the core DAP. Intriguingly, C2CD3, microtubule triplet, and ODF2 jointly constitute the spatial basis of DAP, suggesting a unique configuration of the DAP assembly. Moreover, our results show that the distal-layered ODF2 labeled at the N- and C-terminus construct a fastening unit encircling the microtubule triplets. Together, we develop an organelle-based drift correction protocol and a two-color solution with minimum crosstalk, allowing a robust localization microscopy imaging of expanded cellular structures deep into the gel-specimen composites.

## Introduction

Centriole contributes to multiple critical cellular functions. It plays a vital role in the cell cycle by forming spindle fibers and initiating the cell cycle^1^. The mother centriole is uniquely responsible for ciliogenesis as the basal body of a primary cilium^2–5^. The primary cilium, protruding from the distal end of the basal body, functions as a sensory hub to mediate the transduction of diverse signals^6^. Distal appendages (DAPs) and subdistal appendages (sDAPs) are present at the distal end of the mother centriole. DAPs are shaped as nine-fold symmetric, pinwheel-like structures protruding from the mother centriole. DAPs are essential for ciliary vesicle docking during ciliogenesis^7–10^ to regulate axoneme growth and represent a part of the ciliary gate^11–13^, while sDAPs serve a role in microtubule (MT) anchoring^14^. Numerous DAP proteins have been identified in mammalian cells for their roles in ciliogenesis and ciliary-associated regulations^10, 15–19^ To initiate DAP assembly, C2CD3 is recruited to the centriole distal end^18, 20^, followed by CEP83, which is indispensable for further recruitment of CEP89 and SCLT1. SCLT1 is then required to recruit FBF1 and CEP164^10^. Besides, ODF2, an sDAP-associated protein, has been suggested for its dual function across sDAPs and DAPs^21–23^.

Due to the spatial complexity of appendage structures, investigating such molecular arrangements becomes a significantly challenging task. Superresolution (SR) microscopy, especially single-molecule localization microscopy (SMLM), has been employed for uncovering protein localizations of DAPs and sDAPs with a spatial resolution of ~20 nm ^23–27^. Our previous work used direct stochastic optical reconstruction microscopy (dSTORM) to map the architecture of mammalian DAPs and revealed the novel structure of the DAP matrix between adjacent blades^26^. A further study using correlative SMLM and electron microscopy (EM) demonstrated a close relationship between DAP protein localization and its electron-dense micrograph^27^. However, the resolving power of localization microscopy is still more than an order of magnitude worse than that of EM, which thus results in different structural interpretations between these two scopes. Moreover, CEP83 is reported to own a minimum measure, yet the statistical result still shows a detectable separation of over 50 nm from the centriole wall, implying a missing root structure for the described DAPs^26^. While the superresolution imaging suggested that C2CD3 is concentrated in the centriole lumen^26, 28, 29^, a direct structural relationship between C2CD3 and other core DAP proteins remains elusive due to insufficient technical evidence based on the given mapping precision. These imply that we urgently need an advanced imaging modality with resolution beyond current SR techniques for investigating the missing root configuration of DAPs.

By embedding cellular structures into the network of swellable polyelectrolyte hydrogel, one can physically expand a biological specimen to enable SR imaging under conventional fluorescence microscopy— referred to as expansion microscopy (ExM)^30–32^. Numerous expansion strategies have been proposed for different purposes, for example, magnified visualization of protein, RNA, and transcriptomics in various specimens: cells, isolated centrioles, tissues, and brains^33–37^. Further resolution enhancement can be achieved by integrating ExM with structured illumination microscopy (SIM)^38^, stimulated emission depletion microscopy (STED)^39^, or SMLM^40, 41^. Notably, a combination of ExM and localization microscopy enables us to push the resolution limit of a light microscope well close to an EM level. Nonetheless, the increased distance between organelles and coverslips makes drift correction an arduous task, limiting us to finding proteins of interest only near the coverslip. The deficiency of a pragmatic imaging solution and drift correction protocol also hinders the benefits of multi-color imaging in practice. Although singlemolecule localization imaging of expanded samples ideally demonstrates its ability to resolve the ultrastructural features, general applications to a broad spectrum of biological questions remain challenging.

In this study, we strategically integrate ExM with dSTORM (Ex-dSTORM) to elucidate ultrastructural details of mammalian DAP root structure in intact cells. First, we proposed a practical workflow incorporating in-situ drift correction to enable Ex-dSTORM imaging throughout the entire cell, not limited to coverslips. Second, we optimized the combination of red and far-red dyes for yielding minimum spectral crosstalk in two-color Ex-dSTORM. Therefore, using Ex-dSTORM, we reached unprecedented details in understanding DAPs and gained extensive insight into their three-dimensional (3D) ultra-construction of the higher-order protein complexes. Finally, we uncovered the root structure of DAPs and bridged the spatial relationship between DAPs and centriole to precisely interpret the 3D computational model.

### In-situ drift correction enables systematical sub-5-nm protein mapping with optimized Ex-dSTORM

To systematically investigate the ultrastructural details and protein-protein spatial relationship of DAPs in mammalian cells, we optimized the Ex-dSTORM from several different perspectives (**Fig. 1**). In our experiment, we prepared samples with a post-labeling expansion procedure. It is evident that this sample preparation offers many advantages high labeling density, reduced linkage error, and reduced antibody competition. Moreover, the monomer concentration used in the post-labeling ExM could affect the ultrastructural context of multiprotein complexes^37^. The workflow of post-labeling ExM is shown in **Fig. 1a**. The optimized perfusion concentration in our experiment (1.4% FA and 2% AA in 1x PBS) can achieve a greater expansion factor and, thus, a higher resolution for DAP structure (**Supplementary Fig. 1**). Special note is given to the re-embedding process. We found that the process with materials (10% AA, 0.15% BIS, 0.05% APS and 0.05% TEMED) in only ddH_2_O– rather than in Tris buffer– could hold ~93% of the original size of the expanded hydrogel after reembedding. Overall, we could obtain approximately 4-fold linear expansion after reembedding (**supplementary Fig. 2**).

**Fig. 1.**
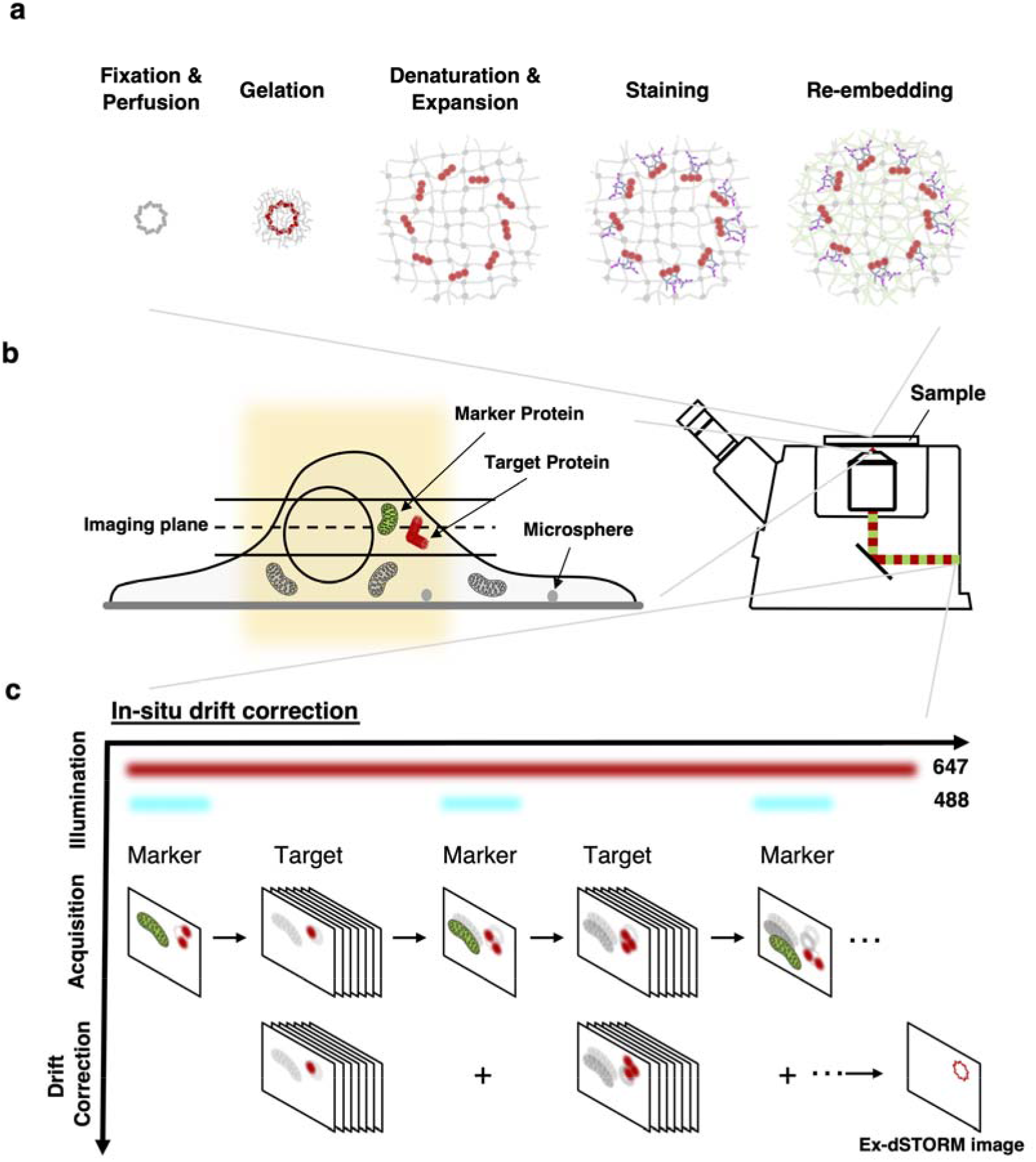
Schematics of optimized Ex-dSTORM with in-situ drift correction. **a** Sample preparation utilizing an optimized post-labeling ExM, including fixation, perfusion, gelation, denaturation, expansion, staining, and re-embedding. Both optimized perfusion concentration and re-embedding solution without Tris were used for a higher expansion factor (details in Methods). **b** In-situ marker labeling enabling in-situ drift correction with Ex-dSTORM imaging. The microsphere beads deposited on the coverslip fail to be the fiducial markers in the expanded sample due to a long distance from the imaging plane at the target protein. **c** Schematic of the in-situ drift correction. Through immunolabeling marker proteins across cells in panel (**b**) (green channel), Ex-dSTORM imaging of target proteins can be performed anywhere inside the expanded gel. Besides the 638-nm excitation light for single-molecule imaging, we intermittently light 488-nm laser to acquire the lateral drift of the sample for postprocessing position compensation. A smooth interpolation between two marker image series is performed owing to slow drifting. Then, the resulting Ex-dSTORM image is obtained by superimposing the localizations of single molecules without marker.

The lateral drift is an inevitable issue in the SMLM because of the long image acquisition, and consequently, a post-processing correction is regularly needed. A specimen expansion procedure usually brings cellular organelles tens of micrometers away from the coverslip. More specifically, fiducial beads (for example, gold beads and microspheres) frequently used in the SMLM are often deposited on the coverslip surface. They are presumably outside the detection focus at the cellular structure of interest, thus causing difficulties in the operation of drift correction (**Fig. 1b**).

Here, we conceived an image-based method for in-situ lateral drift correction by monitoring cellular structures (marker protein) adjacent to the proteins of interest (target protein). During image acquisition of the target protein, we also allow the images of marker proteins to be intermittently recorded for following pattern correlation analysis (**Fig. 1c**). We utilized ATP synthase, for example, as the marker protein, which is enriched and widely distributed over a cell. With the marker feature occupying most of the cell, one can readily locate cells within the expanded gel and then look for the target protein nearby. To our satisfaction, our method no longer constrains the study of cellular structures to merely the coverslip surface. Besides, the intense fluorescence signals from ATP synthase allow samples to endure frequent exposure to the laser light, improving the accuracy of drift correction and hence contributing to high localization precision. Since this approach employs a common detection channel (green) to record marker images, the additional chromatic aberration usually arising from different colors of markers can be avoided. Given that the marker protein is within the hydrogel, any relative movement between markers and targets could be minimized.

Furthermore, we customized a mounting chamber for Ex-dSTORM (**supplementary Fig. 3**). This design provides excellent mechanical stability and ensures that samples are immobilized in the chamber to minimize lateral drift between the cover glass and the sample. Therefore, we only need to correct the drift between the objective and samples. Lastly, we estimated the system resolution of our Ex-dSTORM, which achieves ~3-nm localization precision (**Supplementary Fig. 4**). The resolving power is ideal for investigating the ultrastructure of protein complexes with quantification analysis at a molecular scale.

### Revealing nine-fold symmetry of C2CD3 and ultrastructural contexts of DAP proteins with Ex-dSTORM

Superresolution microscopy has been used to image many centriolar proteins, but the studies on ultrastructural contexts of DAPs have not yet been reported due to insufficient resolution. Hence, we systematically imaged the core DAP proteins (C2CD3, CEP83, CEP89, SCLT1, FBF1, and CEP164) and a centriolar structure protein (Ac-tub) in human retinal pigment epithelial cells (RPE-1). First, we synchronized RPE-1 cells in the G_1_/G_0_ phase to arrest the spatial arrangement of the DAP proteins in the fully-grown state. Then, we conducted Ex-dSTORM imaging of the DAP proteins from the axially oriented mature centrioles by searching for circular ring patterns of protein images. A remarkable improvement in resolution is found in the Ex-dSTORM results compared to other imaging methods (**Fig. 2a**). Notably, the ultrastructural arrangement of DAP proteins at individual blades can only be revealed by Ex-dSTORM. To further determine the organelle-defined expansion factor for a more accurate quantitative analysis, we divided the mean diameter of SCLT1 analyzed from Ex-dSTORM images by that analyzed from dSTORM images (supplementary Table 1). This analysis yields an expansion factor of 3.92, slightly smaller than the value obtained from a direct measurement of gel size. Surprisingly, C2CD3, the core DAP protein essential for DAP assembly, presents as an apparent nine-fold radial symmetry under Ex-dSTORM, which is different from the previous findings that suggested its concentrated distribution in the centriole lumen (**Fig. 2a**). Instead, C2CD3 may show a close spatial correlation with the centriole wall (Ac-tub).

**Fig. 2.**
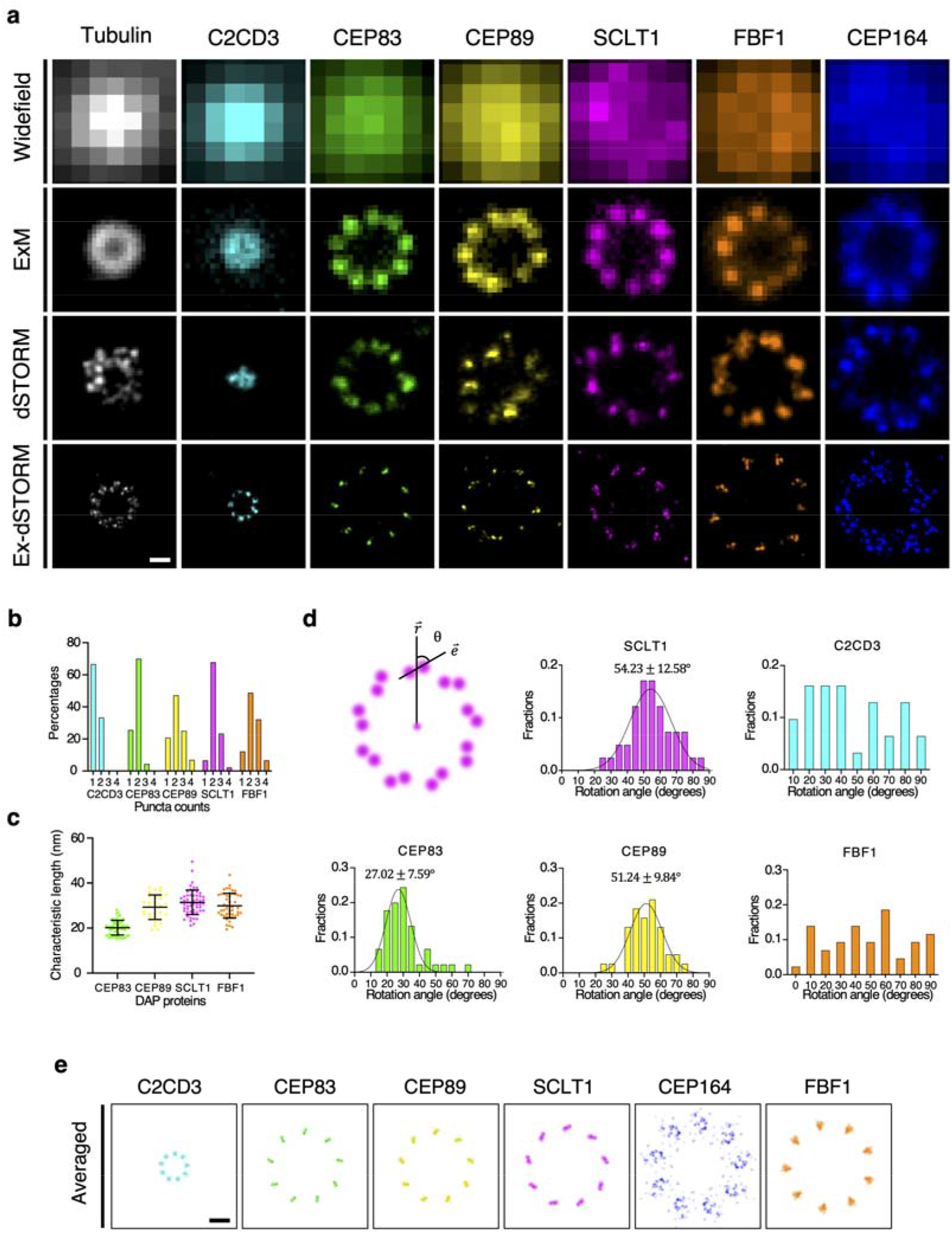
Ultrastructural protein mapping of distal appendage (DAP) achieved by Ex-dSTORM. **a** Representative images of various DAP proteins under different microscopy techniques. With the ~60x resolution enhancement, Ex-dSTORM exhibits ultrastructural details of the DAP proteins, which are not resolvable by other imaging modalities. **b** Histogram analysis of the number of puncta discovered at each blade of DAP proteins. Most DAP proteins show more than one punctum in each blade, forming rod-like patterns (n = 90 for each protein). **c** Characteristic length analysis of DAP proteins with rod-like patterns in (**b**) except C2CD3, which mostly appears as one punctum in a single blade (mean ± SD, n ≥ 31 blades each). This length indicates the 2D axial-view projection of the 3D arrangement of these proteins. **d** Statistical analysis of the rotation angle (*θ*) of DAP proteins with rod-like patterns observed from the distal end. SCLT1, CEP83, and CEP89 exhibit the specific clockwise rotation angles (*θ*) between the radial direction 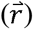 and the characteristic orientation 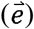, while FBF1 and C2CD3 show a dispersed distribution in angle (n ≥ 31 blades each). **e** Averaged images of DAP proteins enhancing corresponding localization features. In particular, the averaged image of FBF1 demonstrates its possible localizations rather than a specific pattern. Scale bars, 100 nm (**a**, **e**)

For some DAP proteins, our Ex-dSTORM images revealed multiple localization puncta at each blade and organized patterns in these puncta (**Fig. 2 a-d**). Two isolated puncta at individual blades are mainly found for SCLT1, CEP83, and CEP89 (**Fig. 2b**), and their different characteristic lengths (XY-plane projection) may indicate distinct 3D orientations of the DAP proteins (**Fig. 2c**). To determine whether the orientation of puncta is conserved, we also performed statistical analysis by characterizing the angle from the radial direction (**Fig. 2d**). Our results suggest the specific rotation angles of ~27°, ~51°, and ~54° (clockwise) for CEP83, CEP89, and SCLT1, respectively (**Fig. 2d**). Interestingly, these three proteins displayed the same clockwise chirality as MT triplets when viewed from the distal end, while FBF1 and C2CD3 puncta presented a broad angular distribution. As for CEP164, the proteins showed a clear anticlockwise chirality and formed a curved feather-like shape (**Fig. 2a**).

Lastly, we performed a rotational average of the localization signals based on a nine-fold symmetric configuration to feature the ultrastructural orientation among the DAP components (**Supplementary Fig. 5**). The DAP proteins were organized in certain chirality except for FBF1 (**Supplementary Table 2**), implying that FBF1 may involve different ciliary mechanisms other than being a structural component of DAPs (**Fig. 2e**).

### Optimized dye combination for minimum crosstalk in two-color Ex-dSTORM

Spatial relationships among distinct proteins could not be revealed by single-color imaging. Nevertheless, due to several practical challenges, two-color Ex-dSTORM has not been routinely applied to study biological questions. To this end, we optimized two-color Ex-dSTORM with red and far-red channels, which generally demonstrate excellent optical properties for single-molecule signals.

First, Alexa Fluor 647 (AF647), a frequently used dye in two-color SMLM, was used for the far-red channel, and CF568 was used for the red channel. In our experiment, we observed an abundance of additional signals— other than the signals from CF568— in the red channel detection. To this, the observed event is essentially a crosstalk effect. Furthermore, to quantify the crosstalk phenomenon and search for the slightest crosstalk among far-red dyes, we immunolabeled two proteins, C2CD3 with CF568 (which only appeared on the centriole-distal-end, yellow box, **Fig. 3a**) and Ac-Tub with three frequently used far-red dyes, AF647, ATTO647N, and Dyomics 654 (Dy654) (**Fig. 3a**). This protein combination allowed us to recognize crosstalk at the cilium compartment because of their different spatial distribution and protein numbers. In the far-red channel, we only observed the pattern of Ac-Tub under our expectations. Nonetheless, in the red channel, besides the designated pattern of C2CD3 (yellow box, **Fig. 3a**), we also detected the pattern of Ac-Tub (regions marked with blue lines, **Fig. 3a**). We found Dy654 demonstrated the most negligible crosstalk effect in the red channel (below 2% in both analyses) (**Fig. 3b**). On the contrary, given the observation of Ac-tub, AF647 reached over 30% of the crosstalk effect, which may cause significant image artifacts to the red channel. As a result, we chose Dy654 as our far-red dye in the following experiment to provide more reliable information, especially in two-color Ex-SMLM. Hence, this dye combination allows us to accomplish two-color sub-5 nm protein mapping with high fidelity.

**Fig. 3.**
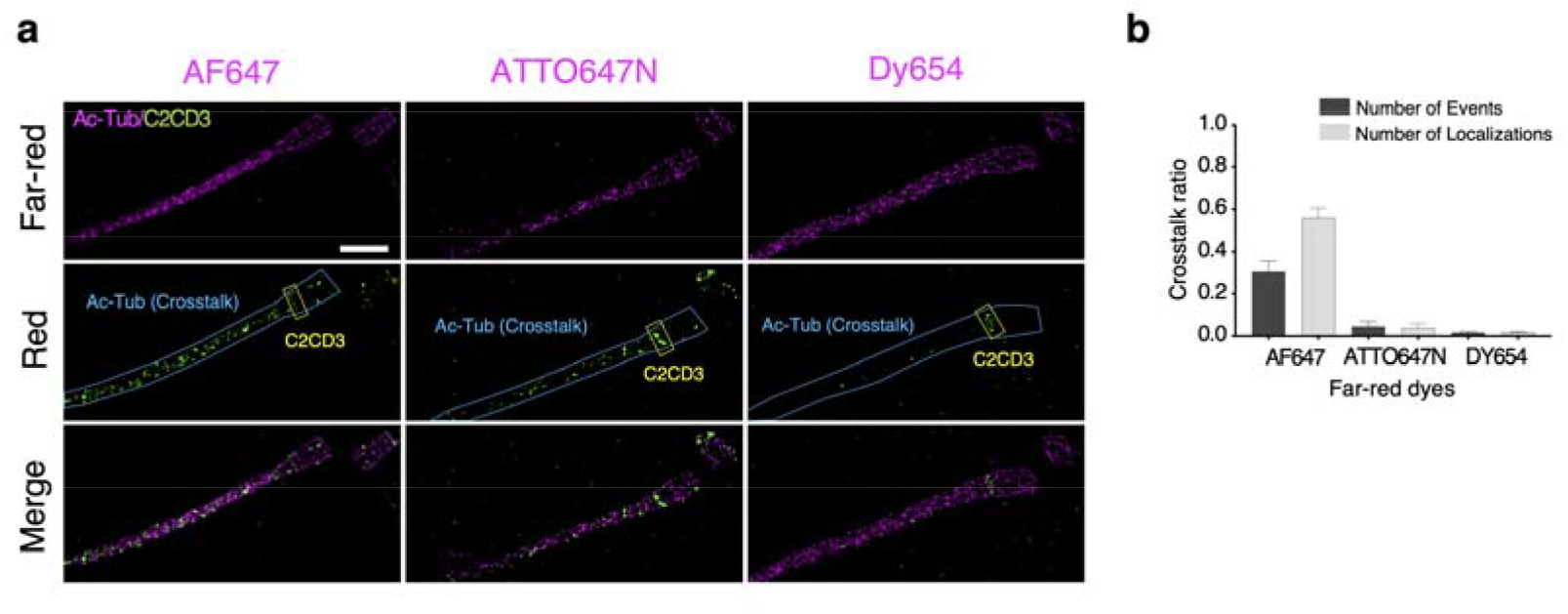
Crosstalk analysis among far-red dyes. **a** Two-color Ex-dSTORM images of C2CD3 labeled with CF568 and Ac-Tub stained with different far-red dyes, displaying a crosstalk effect in the red detection channel. Yellow boxes mark the signals of C2CD3, while blue lines enclose the crosstalk signals from Ac-Tub staining with AF647, ATTO647N, or Dy654. **b** Quantitative analysis of crosstalk ratio among these far-red dyes with two analytical methods based on single-molecule events or localizations. Both results indicate the minimum crosstalk using Dy654 (mean SD, n = 4 for each far-red dye). Scale bar, 500 nm (**a**).

### Ex-dSTORM axial-view imaging reveals ultra-resolved angular and radial relationships among DAP proteins

We next performed minimized-crosstalk two-color Ex-dSTORM imaging from axial orientation to re-define the configuration of DAPs at the molecular level with high fidelity. We have made several discoveries. Firstly, relative allocations of proteins in the radial direction revealed the protein-protein spatial relationships. The proteins C2CD3, SCLT1, CEP83, CEP89, and CEP164 were imaged and paired. From our result, the two-color image pairs of C2CD3-SCLT1, CEP83-SCLT1, CEP83-CEP89, and CEP164-SCLT1 demonstrated that they were all arranged in the same radial direction (**Fig. 4a**-**c**, **e**). By contrast, the image pair of FBF1-SCLT1 showed that FBF1 was in a different radial direction among DAPs (**Fig. 4d**). Moreover, we found that CEP83 partially overlapped with CEP89 and that, statistically, CEP83 displayed a slightly smaller radius (**Fig. 4c**). We also found that SCLT1 was nearly at the centroid of CEP164 patterns and enclosed by the anticlockwise feather-like signals of CEP164 (**Fig. 4e**).

**Fig. 4.**
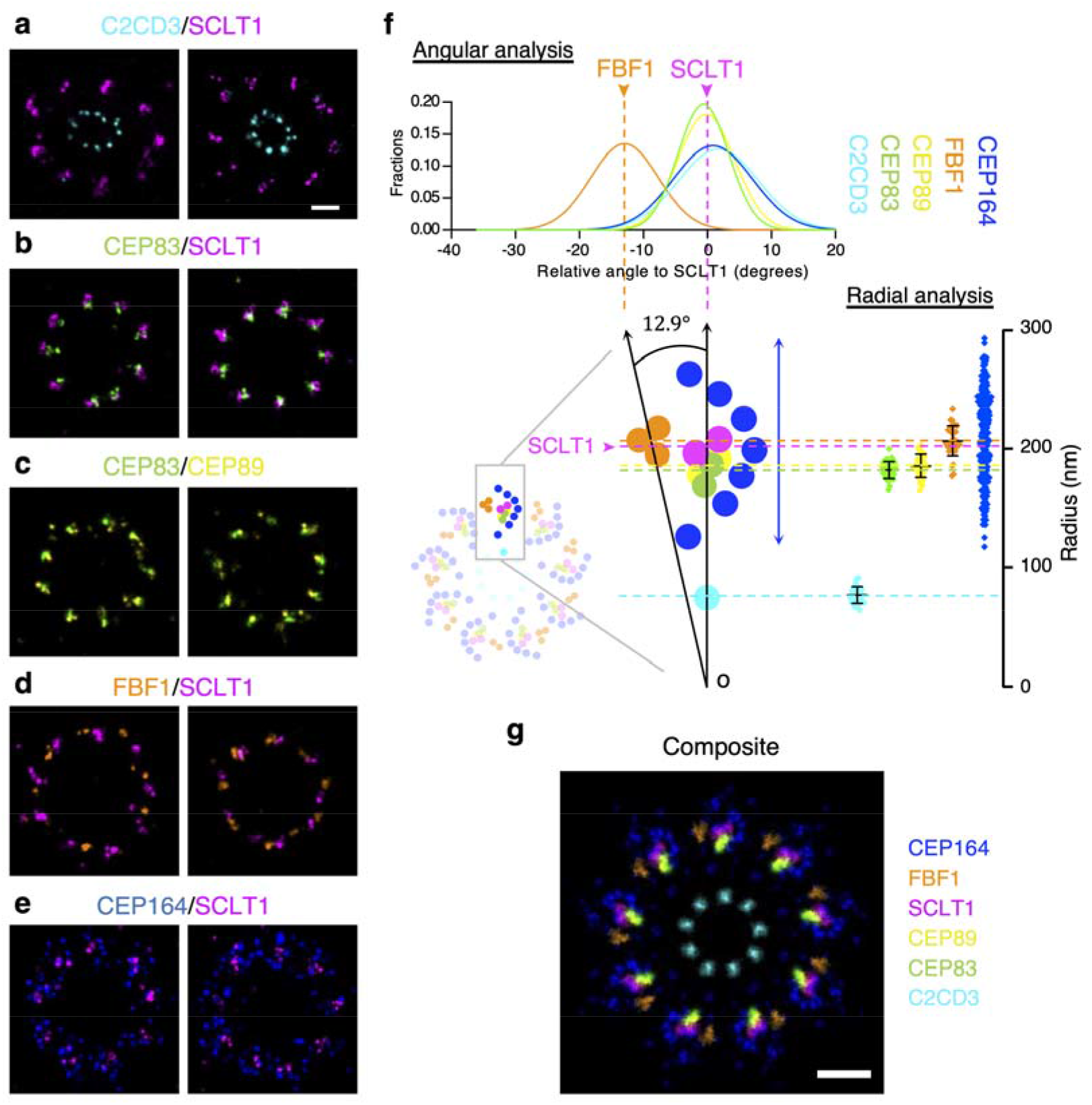
Revealing ultra-resolved angular and radial relationship among DAP proteins with two-color Ex-dSTORM imaging. **a**-**e** Two representative two-color axial-view Ex-dSTORM images. C2CD3-SCLT1 (**a**), CEP83-SCLT1 (**b**), CEP83-CEP89 (**c**), and CEP164-SCLT1 (**e**) pairs align in the same radial direction, whereas FBF1 exhibits a significantly distinct angular position with respect to SCLT1 (**d**). SCLT1 is enclosed by the “feather-like” CEP164 distribution (**e**). **f** Angular and radial analyses of DAP proteins. Angular analysis demonstrates that C2CD3, CEP83, CEP89, and SCLT1 allocate at a concentrated angular distribution around 0° (n = 243 points for each pair), whereas the mean position of FBF1 deviates from SCLT1 by - 12.9° Radial analysis reveals the mean radius of C2CD3, CEP83, CEP89, SCLT1, and FBF1 (mean ± SD, n = 36 blades for each pair), while CEP164 proteins are represented with their individual localizations (n = 261 points from 4 centrioles). **g** Composite of averaged images of DAP proteins (Fig. 2e) scaled with their mean angular and radial positions. FBF1 is exceptionally located in a different radial direction in contrast to CEP83, CEP89, and SCLT1 enclosed by CEP164. Scale bars, 100 nm (**a**-**e**, **g**).

Secondly, we performed quantitative analyses from the axial orientation to unveil the protein-protein angular relationships— angular spacing and relative angle. For starters, the angular spacing of all protein pairs was measured and found to be precisely repeated every 40° interval (**Supplementary Figs. 6 and 7**). Notably, histograms of angular spacing from our Ex-dSTORM images demonstrated a much more satisfactory angular resolution of about 0.5°, crucial for differentiating spatial arrangement among DAP proteins at a few nanometers. Besides angular spacing, we explored relative angles from the histogram analyses using SCLT1 as the reference. The centers of C2CD3, CEP83, CEP89, and CEP164 were arranged at a concentrated angle of around 0° but with distinct distributions (**Fig. 4f**; **Supplementary Fig. 8**). As for the relative angle of FBF1, on the contrary, we found that it localized at a distinctive angle of ~12.9° with respect to SCLT1, suggesting that it was not aligned with other DAP proteins. Nevertheless, regular SMLM could have difficulty differentiating between FBF1 and SCLT1 as it does not have enough resolving power^27^. More importantly, this result supports our previous finding that FBF1 may serve a unique role in ciliary gating in addition to DAP formation ^26^.

Thirdly, we further conducted the radial analysis to investigate the configuration of DAPs. In our result, the radial analysis of C2CD3 revealed neither the disc-like pattern nor concentration in the lumen^28, 29^ (**Fig. 4f**); this suggests that our Ex-dSTORM can outperform typical dSTORM to resolve the nine-fold radial pattern. Besides, it is worth mentioning that we eliminate the cell-to-cell difference and blade-to-blade variation for the radial analysis by measuring the individual radius ratio blade by blade (**Supplementary Fig. 9**).

Taken together, we reconstructed the ultrastructural composite image by assembling the information of radius ratio, relative angle, and averaged images (**Fig. 4g**). Surprisingly, although most DAP proteins appeared in the clockwise direction, the resulting profile in the composite represented the anticlockwise direction.

### Ultra-detailed analyses of the relative longitudinal relationship among DAP proteins

Using SCLT1 as the reference, we obtained a relative longitudinal position among DAP proteins. Our result denoted a few intriguing features of these DAP proteins from the lateral view with a series of two-color Ex-dSTORM imaging. First, CEP83 possessed two-layered populations separating further in the longitudinal direction; moreover, CEP83 constructed a colinear correlation with SCLT1 (gray lines, **Fig. 5a**). CEP89, SCLT1, and FBF1, in contrast to CEP83, mainly exhibited nearly horizontal distributions and formed a certain inclination angle (dashed lines, insets, **Fig. 5a**). Second, to underline the stereoscopic sense of DAPs structure, we further captured two-color Ex-dSTORM images from a tilted view. A combination of CEP83-SCLT1 revealed the crown-like configuration (tilted view, **Fig. 5a**). While CEP83-CEP89 showed similar values in radius, they manifested an explicit offset in the longitudinal orientation compared to the axial observation (tilted view, **Fig. 5a**, **Fig. 4c**). Third, we found that most inner molecular localizations of CEP164 presented at lower longitudinal positions (arrowheads, **Fig. 5a**). In addition, we confirmed that SCLT1 was located within the geometric center of CEP164 as if a curve feather of CEP164 was embracing SCLT1 (tilted view, **Fig. 5a**).

**Fig. 5.**
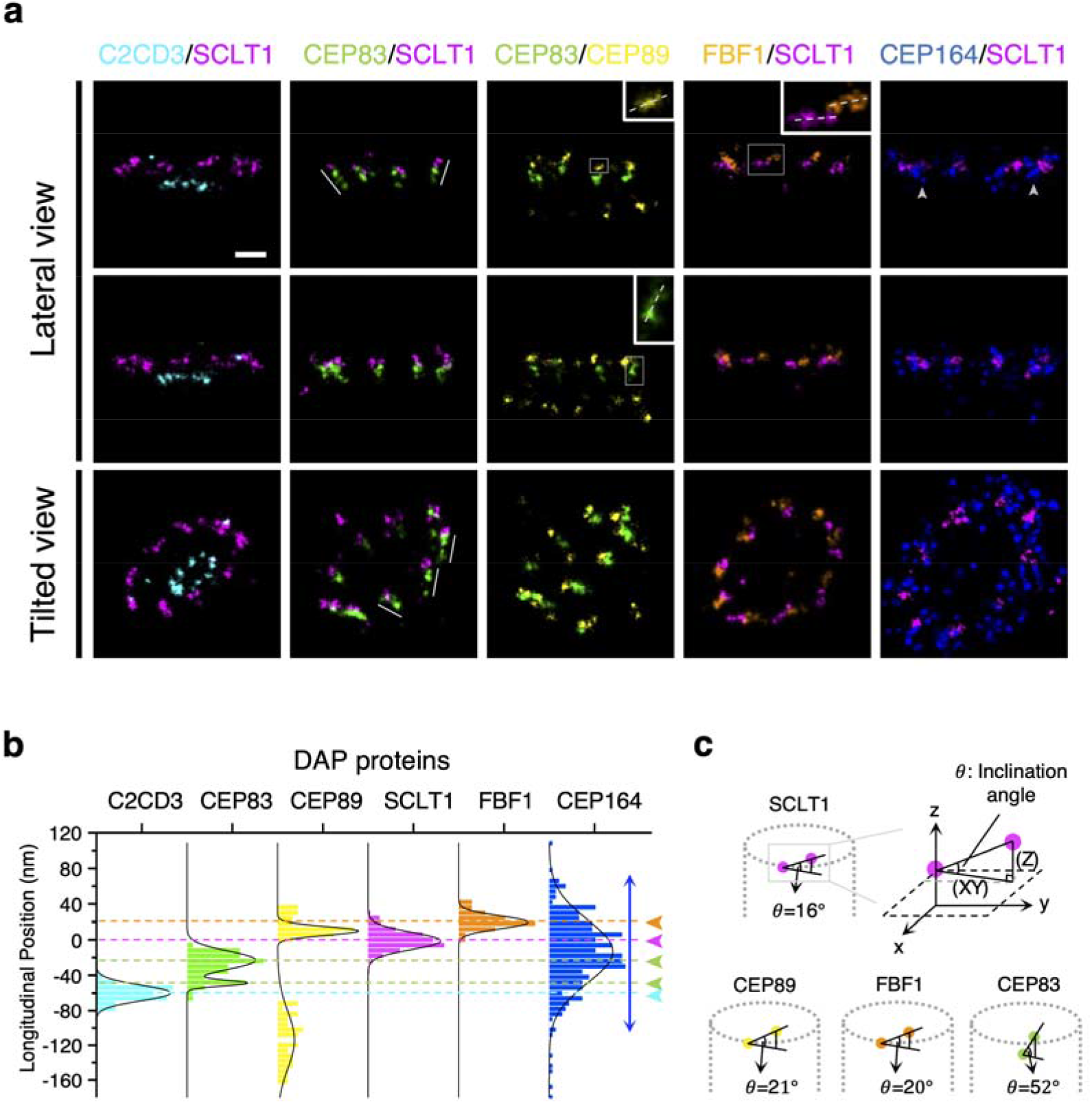
Lateral-view Ex-dSTORM images enabling the ultrastructural analysis of the longitudinal position of DAPs. **a** Representative two-color Ex-dSTORM images from lateral view and tilted view. Each pair of DAP proteins from lateral-view images illustrates their relative longitudinal positions. The collinear relationship is particularly found in the CEP83-SCLT1 images (gray lines). The angle of inclination of each DAP protein is identified for CEP83, CEP89, FBF1, and SCLT1 (dashed lines, insets). Gray arrowheads mark the narrower radial distributions at the lower bounds of CEP164 longitudinal localizations. The tilted-view images highlight the stereoscopic interpretation of the spatial arrangement of DAP proteins. **b** Histogram analysis of the longitudinal positions of DAP proteins relative to SCLT1 (n ≥ 7 centrioles for each). Arrowheads indicate the mean longitudinal position of each DAP protein except for CEP164, which distributes throughout the other proteins (blue double-headed arrow). **c** Computation of the inclination angle of DAP proteins at each blade for SCLT1, CEP89, FBF1, and CEP83 by the ratio of the longitudinal projection (Z) to axial projection (XY). Scale bar, 100 nm (**a**).

To quantify the longitudinal position of each DAP protein, we again used SCLT1 as the reference to determine each localization of molecular puncta. Ex-dSTORM imaging enabled us to perform the paired measurement in both the axial and lateral view (**Supplementary Fig. 9**). Therefore, we could rule out the longitudinal position variation from different blades, obtaining an accurate molecular position of each protein, not merely reporting an average position (**Fig. 5b**). Histogram analysis disclosed the two-layered CEP83 distribution, which was separated by ~ 26.0 nm (green arrowheads, **Fig. 5b**). The histogram analysis also revealed a broad distribution of CEP164 along the longitudinal direction (blue arrows, **Fig. 5b**). For molecular orientation, the inclination angle of CEP83, CEP89, SCLT1, and FBF1 with ~ 52.1°, ~ 21.0 °, ~ 16.4°, and ~ 19.5° can be determined from axial (XY) and lateral (Z) projection (**Fig. 5c**; Supplementary Table 2). In particular, we found that the inclination angle of CEP83 is in good agreement with that of the DAP blades from the EM micrographs ^24, 27^-this indicated that CEP83 represented a direct structural involvement in the DAP profile.

### Ex-dSTORM identifies a missing coverage over the DAP root structure

Previous studies frequently overlapped SMLM with EM images to bridge the spatial relation between DAP and centriole^24, 26, 27^. Nonetheless, resolutions between the two techniques diverge an order of magnitude apart. Hence, it is prone to different data interpretations without comparable image quality between different methods^26, 27^. Moreover, the root DAP structure near the centriole was unclear due to an insufficient resolution. In order to directly and precisely correlate DAP structure with a centriole, we first imaged Ac-Tub to examine centriolar triplets, which displayed more clearly in the averaged image (**Supplementary Fig. 10**). Next, we performed two-color Ex-dSTORM imaging of Ac-Tub and C2CD3. Interestingly, our result shows that C2CD3 localized between adjacent triplets, not concentrating in the centriole lumen and that C2CD3 was arranged in an extension line of triplets (gray lines, **Fig. 6a**). To determine whether C2CD3 is close to A-tubule or C-tubule end, we quantified the correlation of C2CD3 and Ac-Tub with radial and angular information. We first evaluated the radius ratio of C2CD3 over MT triplets, and the value of 0.733 was obtained (**Fig. 6b**). To precisely determine the angular relationship between C2CD3 and Ac-Tub, we measured the angle between the center of the triplets and their adjacent C2CD3. Our finding showed that C2CD3 proteins were positioned between triplets near A-tubule (θ ~ 20.5°, **Fig. 6b**). Incorporating the information of DAP proteins/C2CD3/Ac-Tub, we illustrated the coverage of these proteins in their radial distributions (**Fig. 6c**). Notably, because of significant improvement in resolution and accurate labeling sites with Ex-dSTORM, we found a gap between CEP83 and Ac-tub-which implies a missing part of the DAP root structure.

**Fig. 6.**
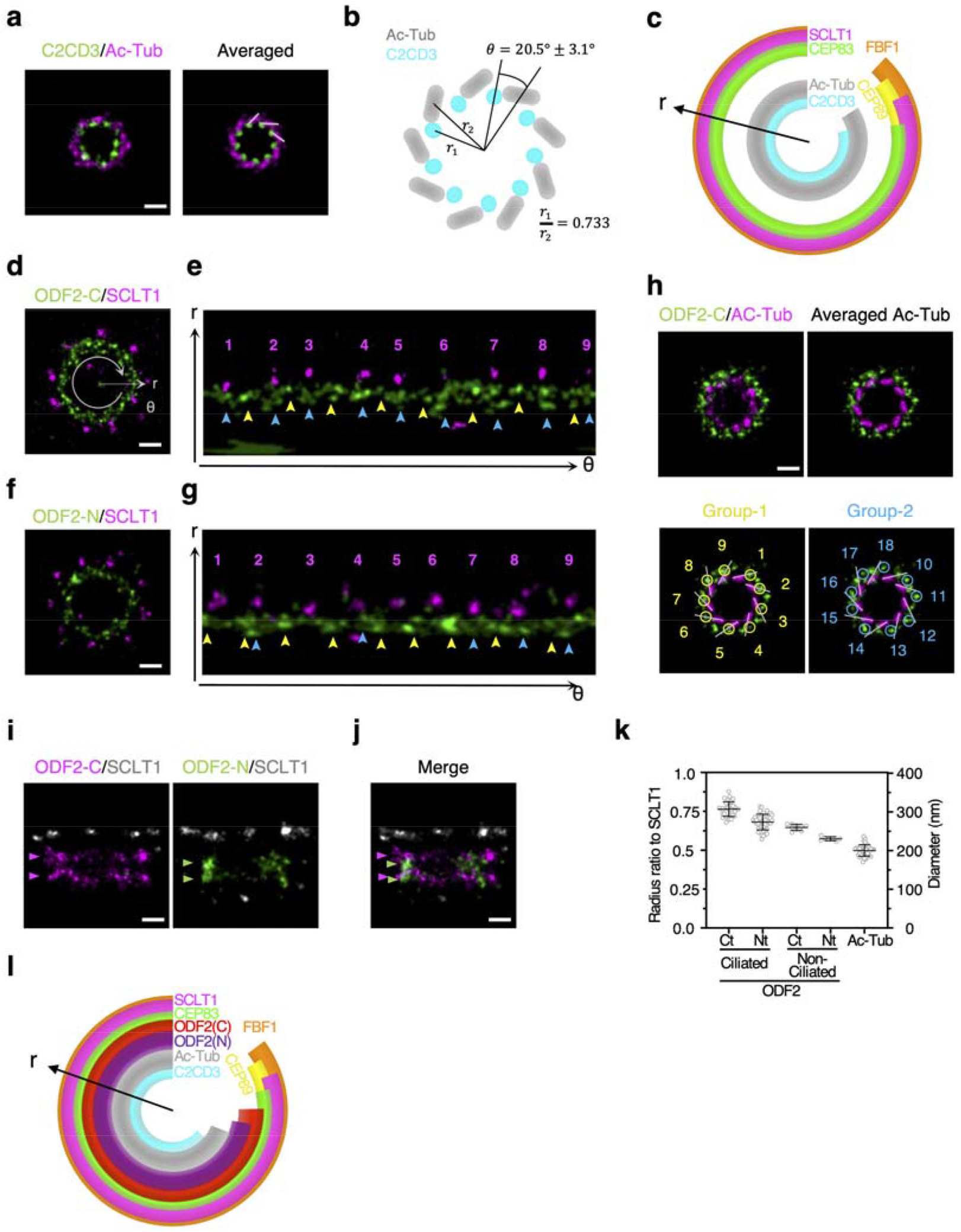
Spatial correlation between DAP and centriole revealing the root structure of DAPs. **a** A representative two-color Ex-dSTORM image of C2CD3/Ac-Tub disclosing the direct spatial relationship between MT triplets and DAP. The image enhancement is further achieved by rotational averaging every 40°. The result shows that C2CD3 localizes inwards at the extension line of Ac-Tub (gray lines). **b** Illustration of the spatial relationship between C2CD3 and Ac-Tub denoting their angular difference and radius ratio (n = 42 blades for radial analysis, n = 5 centrioles for angular analysis). **c** Radial coverage of DAP proteins and MT triplets implying a missing root structure of DAP (mean ± 3 SD for each protein). **d**, **f** Representative two-color Ex-dSTORM images in the axial view of the C-terminus of ODF2 (**d**) and the N-terminus of ODF2 (**f**) with SCLT1. **e**, **g** Polar unwrapping analysis revealing that both ODF2-C and ODF2-N can be characterized into two groups-one locating at the same angular position (*θ*) with SCLT1 (blue arrows) and the other localizing between SCLT1 (yellow arrows). **h** Two-color Ex-dSTORM image of ODF2-C/Ac-Tub manifesting two-grouped populations of ODF2. Again, the Ac-Tub signals are enhanced by rotational averaging. Group-1 of ODF2-C locates at the extension line of Ac-Tub signals (yellow circles). Group-2 of ODF2-C locates between two adjacent MT triplets (blue circles). **i** Representative two-color Ex-dSTORM images in lateral view of ODF2-C and ODF2-N with SCLT1. Two-layered distribution can be clearly distinguished in both ODF2-C and ODF2-N. **j** Merged image from (**i**) for comparing longitudinal position. **k** Quantitative analysis of ODF2-C/N radii for ciliated and non-ciliated cells. Ac-Tub is also compared to non-ciliated ODF2-C/N. **l** ODF2-C/N as the root structure filling the coverage gap in panel (**c**). Scale bars, 100 nm (**a**, **d**, **f**, **h**, **i**, **j**)

To examine how the DAP base was organized, we further characterized ODF2, an sDAP component, as the possible DAP root structure because of its intimate relation with DAP in terms of its spatial position or biological function^21–23^. To elucidate the configuration of root DAP, we immunostained ODF2 at its C- and N-terminus, respectively. Although two-color Ex-dSTORM images of ODF2-C/SCLT1 and ODF2-N/SCLT1 demonstrated a continuous distribution rather than a nine-fold radial pattern as other DAP proteins (**Fig. 6d** and **f**), we could still classify ODF2-C and ODF2-N into two different groups of distributions by polar unwrapping analysis. Our result revealed that one group of puncta patterns aligned with SCLT1 with the same angular direction (blue arrows, **Fig.6e** and **g**); another group was positioned between two adjacent SCLT1 (yellow arrows, **Fig. 6e** and **g**). To observe the pattern of ODF2 clearly, we further imaged ODF2-C with Ac-Tub from the axial orientation to scrutinize its spatial arrangement with a different reference (**Fig. 6h**).

Interestingly, two groups of arrangement could be recognized clearly in the ODF2-C with averaged Ac-Tub: one is distributed at the extension lines of MT triplets (yellow circles, Group 1, **Fig. 6h**), while the other is localized between two extension lines of MT triplets (blue circles, Group 2, **Fig. 6h**). We then move forward to image ODF2-C/N from lateral orientation to elucidate the difference between these two. The two-color Ex-dSTORM images from the lateral view revealed that both ODF2-C/N exhibited a two-layered distribution (arrows, **Fig. 6i**). Moreover, aligning with SCLT1, we found that ODF2-C/N form alternating two-layered distribution beginning with distal ODF2-C layer followed by distal ODF2-N layer, proximal ODF2-C layer, and proximal ODF2-N layer (**Fig. 6j**). Assembling the information of two groups of ODF2-C/N from axial and lateral views, we inferred that one group of ODF2-C/N from the axial view belongs to the upper layer and the other belongs to the lower layer.

Intriguingly, the quantitative analysis of the ODF2-C/N radii showed that the ODF2 radius of the non-ciliated centrioles was systematically smaller than that of the ciliated cells (**Fig. 6k**), indicating the ciliation-dependent distribution for ODF2. Given the result from the radial analysis, our finding suggests that ODF2 fills the undefined coverage, possibly participating in being the root structure of DAPs (**Fig. 6l**).

### Ex-dSTORM unravels the distal-layered ODF2 as the root structure of DAP

Longitudinal position analysis demonstrated that the distal-layered ODF2-C/N partially overlapped with DAPs (**Supplementary Fig. 11**), while the proximal-layered ODF2-C/N was in the region of sDAPs. Moreover, we found a ~18° offset between the aforementioned two groups of ODF2 (**Supplementary Fig. 12**). Hence, we assumed the distal-layered ODF2-C/N to be the root structure of DAPs and the proximal-layered ODF2-C/N to be a part of sDAPs. DAPs and sDAP may form in a staggered arrangement stemming from the angle difference between the two groups. To test this hypothesis, we imaged the sDAP-depleted centrioles by studying CEP128 CRISPR/Cas9 knockout cells^23^ to investigate the ODF2 localizations under Ex-dSTORM. Astonishingly, unlike the two-group distributions in WT cells, a canonically nine-fold symmetric pattern re-emerged in both ODF2-C and ODF2-N images in the sDAP-deficient cells (arrows, **Fig. 7a** and **b**).

**Fig. 7.**
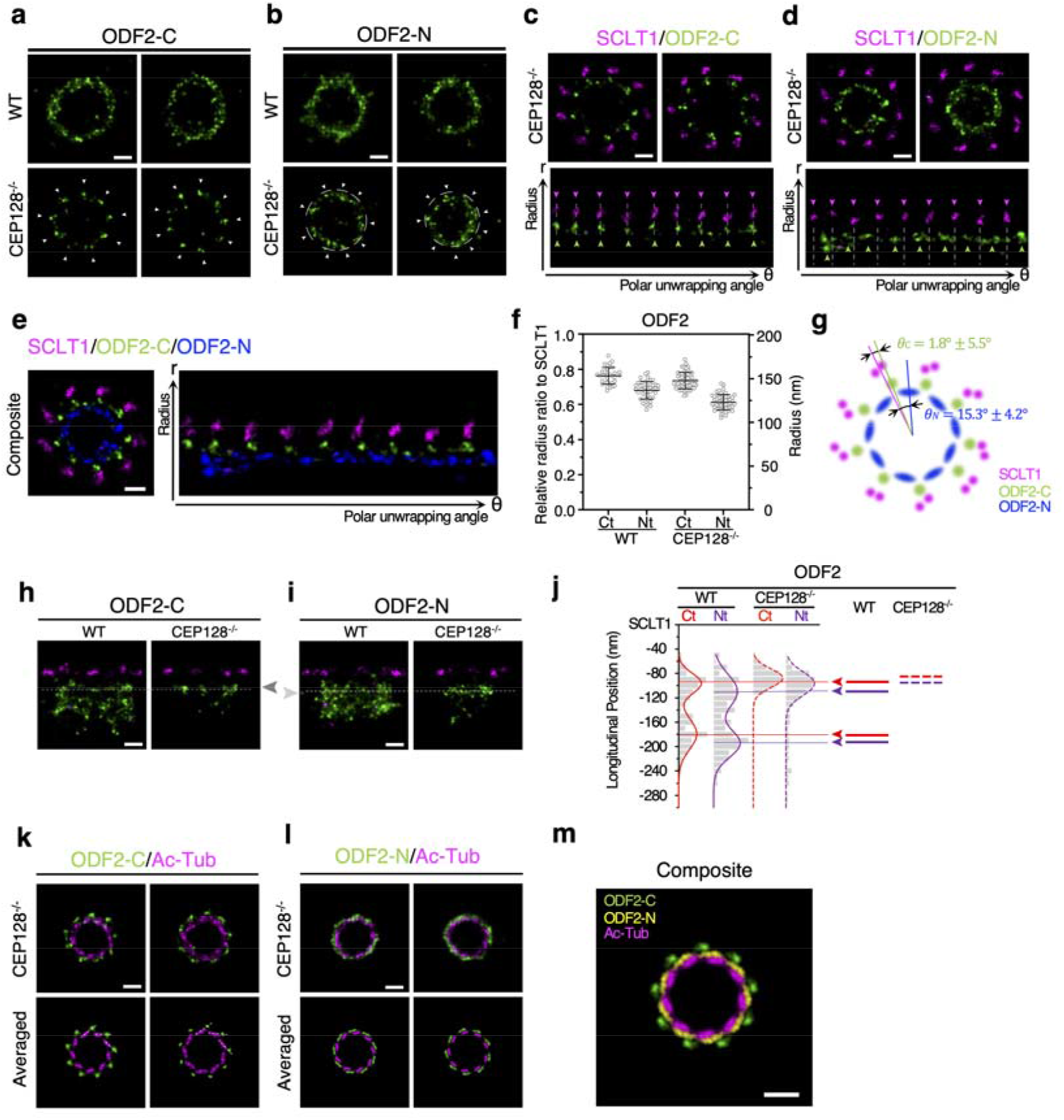
Two-color Ex-dSTORM unraveling the distal-layered ODF2 as the root structure of DAPs. **a**, **b** Representative Ex-dSTORM axial-view images of the C-terminus (**a**) and the N-terminus (**b**) of ODF2 in WT and CEP128^-/-^ cells. The well-resolved nine-fold radial symmetry re-emerged in sDAP-depleted (CEP128^-/-^) cells (white arrows). **c**, **d** Two-color Ex-dSTORM images allowing for differentiation of distinct patterns of ODF2-C/N with SCLT1 in CEP128^-/-^ cells. Polar unwrapping analysis in (**c**) reveals that ODF2-C aligns well with SCLT1, while ODF2-N in (**d**) locates between two SCLT1 puncta. **e** Composite images of (**c**) and (**d**) aligned with SCLT1. Special note is given to the interlacing spatial relationship between ODF2-C and ODF2-N. **f** Mean radius analysis revealing differences of ODF2-C/N between WT and CEP128^-/-^ cells (mean ± SD, n ≥ 36 blades for each condition) **g** Schematic model of ODF2-C/N and SCLT1 with quantitative analysis in angle (mean ± SD, n = 6 centrioles each). **h**, **i** Representative Ex-dSTORM lateral-view images of the C-terminus (**h**) and the N-terminus (**i**) of ODF2 in WT and CEP128^-/-^ cells. The remaining layer of ODF2 in CEP128^-/-^ cells presents as the distal layer in WT cells (dashed line). **j** Longitudinal distribution histogram describing ODF2-C/N relative to SCLT1 for WT and CEP128^-/-^ cells. In both cells, ODF2-C is slightly closer to SCLT1. Upon sDAP depletion, ODF2 is seen moving toward SCLT1 (n ≥ 5 centrioles each condition). **k**, **l** ODF2/Ac-Tub Ex-dSTORM images in CEP128^-/-^ cells. Averaged images highlight the significantly different arrangement ODF2-C locates on the extension line of Ac-Tub (dashed lines) (**k**) while ODF2-N distributes at the periphery of MT triplets (**l**). **m** Composite image from (**k**, **l**) revealing a unique architecture formed by ODF2-C, ODF2-N, and Ac-Tub. Scale bars, 100 nm (**a**-**e**, **h**, **i**, **k**-**m**)

The relationship between the remaining layer of ODF2 and DAP was then characterized by two-color Ex-dSTORM (**Fig. 7c** and **d**). Strikingly, ODF2-C was allocated in line with SCLT1 (**Fig. 7c**). Contrarily, ODF2-N was located between the two adjacent SCLT1 (**Fig. 7d**). In the composite image (**Fig. 7e**), the radius of ODF2-C is shown to be larger than that of ODF2-N. Moreover, ODF2-C/N interlaced with each other, while ODF2-N manifested a broader distribution in the circumferential direction. Later, with quantitative analysis of the radius ratio in WT and CEP128^-/-^ cells, we found a systematically smaller radius ratio of ODF2-C/N in CEP128^-/-^ cells (**Fig. 7f**). This result indicates that the morphology of the DAP would alter upon sDAP depletion. A schematic model for positioning ODF2-C/N and SCLT1 viewed from the distal end is illustrated in **Fig. 7g** with quantitative data.

To determine how the remaining layer of ODF2-C/N in sDAP-depleted cells is associated with that in the WT cells, we utilize SCLT1 as the reference to analyze the longitudinal position of ODF2-C/N with two-color Ex-dSTORM. Our images showed that the remaining layers of ODF2-C/N in CEP128^-/-^ cells precisely aligned with the distal layer in WT cells, implying the role of distal-layered ODF2 as the DAP root structure (**Fig. 7h** and **i**). Histogram analysis verified this result, except that the ODF2-C/N slightly moved toward SCLT1 (**Fig. 7j**). In addition, to explore the spatial relationship between distal-layered ODF2 and MT triplets, we performed two-color Ex-dSTORM imaging of the ODF2-C/Ac-Tub (**Fig. 7k**). Our findings showed that ODF2-C was localized on the extension line outside the triplets (dashed lines, **Fig. 7k**), representing group 1 of ODF2-C (**Fig. 6h**). This finding illustrates that group 2 of ODF2-C refers explicitly to the proximal-layered ODF2-C that supports the formation of sDAPs. By contrast, the images of the ODF2-N/Ac-Tub exhibit that the ODF2-N are distributed at the periphery of MT triplets (**Fig. 7l**). Together, we have constructed the composite image of these three proteins to uncover the relationships among ODF2-C, ODF2-N, and MT triplets (**Fig. 7m**).

## Discussion

In summary, by utilizing optimized two-color Ex-dSTORM, we have demonstrated the power of ~3 nm protein mapping in unraveling the root structure of centriolar DAP, vital for functional and ultrastructural studies. In this work, we proposed the insitu drift correction protocol allowing us to observe the organelles inside the expanded cells through dSTORM. Systematical two-color Ex-dSTORM imaging from the axial and lateral views enables us to deduce detailed 3D molecular arrangement of DAP proteins. The direct relationship between DAPs and MT triplets can be discovered at a few-nanometer resolution. Furthermore, we have identified a missing part of the DAP at its root based on the radial coverage. Next, we have shown that the distal-layered ODF2 acts as the root structure of the DAP (**Fig. 8a**). Integrating the information from the various features (chirality, inclination angle, characteristic length, puncta count, mean position) of each DAP protein (Supplementary Table 2-4), we build a 3D computational model for seeing the single-blade ultrastructure (**Fig. 8b**-**f**). Our results manifest the nearly same radial direction among C2CD3, ODF2-C, CEP83, CEP89, and SCLT1 in the 3D model (dashed line, **Fig. 8b**). Together with the genetic relationship of C2CD3 as an upstream protein in DAP assembly^18, 20, 28, 29, 42^, our model shows that C2CD3 near the A-tubule of an MT triplet is involved in the configuration of the DAP assembly. Presumably, DAP blades are developed through a directional correlation of the following proteins: C2CD3, MT triplets, ODF2-N, ODF2-C, CEP83, CEP89, SCLT1, and CEP164. The distinct angular position of FBF1 with disorganized patterns implies that it may not be a part of the backbone of the DAP blade, and it substantiates the statement that FBF1 performs an essential role in ciliary gating^26^. The 3D model depicts the stereoscopic arrangement of a feather-shaped CEP164 embracing CEP83, CEP89, and SCLT1 (**Fig. 8c** and **d**). ODF2-N encircles MT triplets with broader distributions along the circumferential direction rather than appearing in the same radial direction as other core DAP proteins. This arrangement has more contact sites and may provide structurally higher mechanical stability for DAP formation. In the lateral view, ODF2-C and ODF2-N may jointly construct the pedestal of the core DAP proteins (**Fig. 8e**). A single blade of the 3D model delineates the molecular constitution of DAP from the centriole wall, manifesting our optics-based ultrastructural protein mapping (**Fig. 8f**).

**Fig 8.**
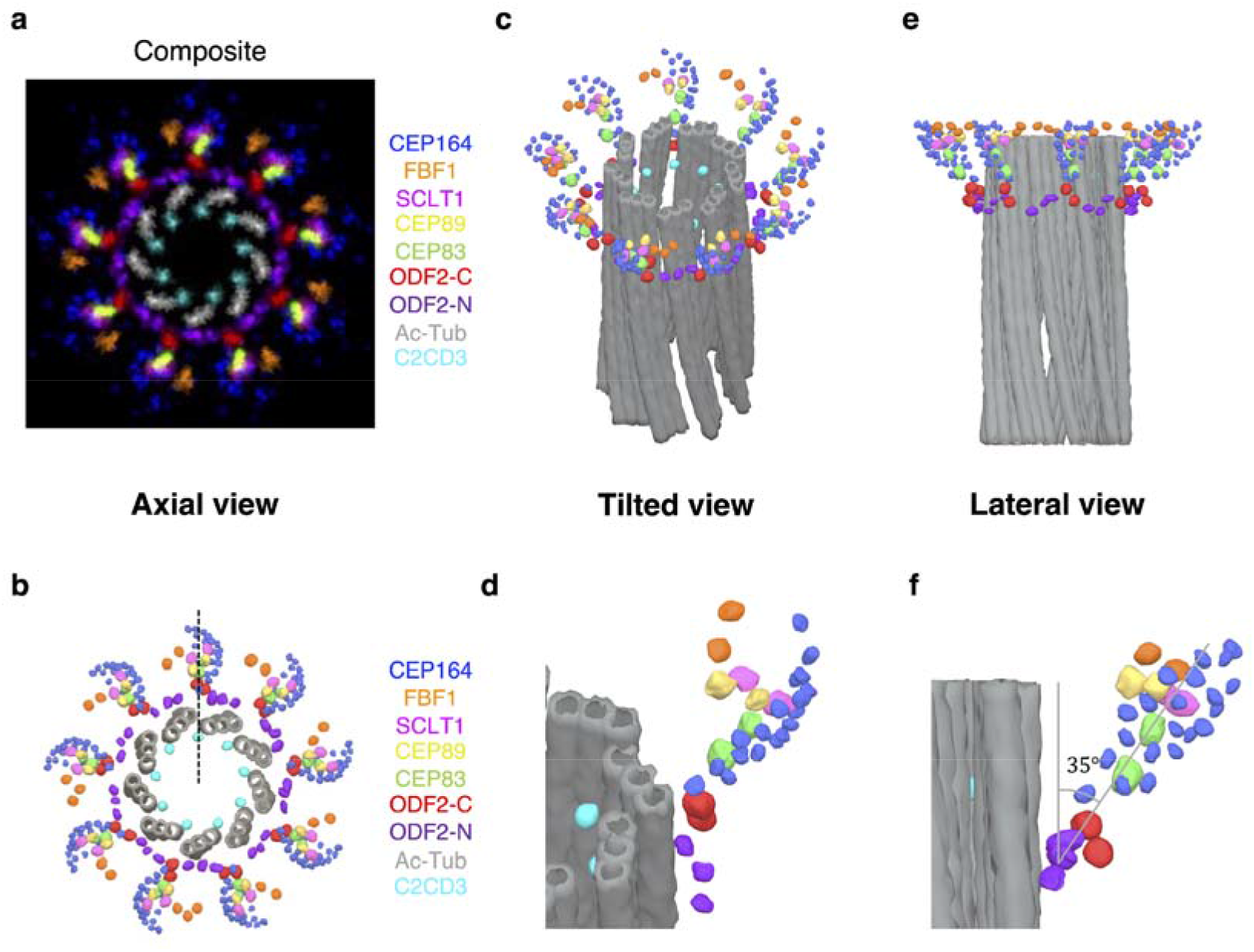
3D model of DAPs displaying the ultrastructural framework from the centriole wall. **a** Composite of rotationally averaged images of the DAP proteins (from **Fig. 1e**) with Ac-Tub (from Supplementary Figure 10) and one example image of ODF2-C/N with scaled dimensions relative to SCLT1. Distal-layered ODF2-C/N fill the aforementioned missing coverage in a root structure of DAP from axial view. **b** Axial-view 3D model positioning individual localizations of DAP proteins, distal-layered ODF2-C/N, and MT triplets (viewed from the distal end of the centriole). The model reveals the same radial direction among C2CD3, ODF2-C, CEP83, CEP89, and SCLT1 (dashed line). The overall configuration of proteins outside the centriole manifests a pinwheel-like morphology revealed by the EM study^49^ **c**, **d** A tilted view of the 3D model illustrating the molecular configuration of all proteins against the mother centriole. Compared with ODF2-C, ODF2-N surrounds MT triplets with a broader distribution along the circumferential direction. A magnified view (**d**) of a single blade manifesting a stereoscopic, feather-shaped arrangement of CEP164, which uniquely embraces CEP83, CEP89, and SCLT1. **e** A lateral view of the 3D model revealing the relative longitudinal positions of all proteins against the mother centriole. ODF2-C/N construct the pedestal of the core DAP proteins. **f** An enlarged view of a single blade in (**e**) delineating that a DAP blade is initiated from the centriole wall, scaffolded via ODF2, and extended to other DAP components. Quantitative measurement of the protrusion angle is highly in accord with previous EM analyses^24, 27^. Scale bar, 100 nm (**a**)

Despite that the previous studies indicated that ODF2 was not directly responsible for DAP formation in RPE-1 cells^10, 43, 44^, our Ex-dSTORM images uncovered the DAP root structure, which was correlated explicitly with the distal-layered ODF2. Hence, the distal-layered ODF2 may act as an auxiliary or indirect role for complete DAP structural construction. Furthermore, the ~18° offset between two layers of ODF2 could cause a corresponding angle difference between DAPs and sDAPs to initiate the specific developing manner of DAPs and sDAPs.

To perform the Ex-dSTORM imaging of a specific organelle, one should first choose a preferred scheme in sample expansion. Although this work has demonstrated a successful application to the study of distal appendages, further procedure compatible with other cellular structures remains to be optimized^45^. Second, we have developed the in-situ drift correction to enable cell imaging deep into a gel and facilitate cell search in various cellular structures. This implementation offers a robust and straightforward solution in most localization microscopy systems.

On the other hand, the crosstalk from the far-red to red channel has posed severe image artifacts in two-color SMLM, which may stem from the photoblueing effect suggested by a recent study^46^. Nonetheless, our result, AF647 causing high crosstalk, does not agree with the previous finding, reporting a minor blue-shifted photoconversion in the oxygen-scavenging system. Perhaps, this contradiction implies more factors to be considered for a holistic understanding of the crosstalk effect in multi-color SMLM imaging. The conclusion in **Fig. 2** was drawn based on considerably different protein numbers-C2CD3 and Ac-tub. In reality, with a proper labeling strategy in far-red and red channels (for example, more proteins for the red channel; fewer proteins for far-red), one can still obtain two-color Ex-dSTORM images with negligible crosstalk (**Supplementary Fig. 13**).

The emerging technique of synergistically combining expansion microscopy and single-molecule localization microscopy sheds light on studying protein complexes at the molecular level. However, practical localization microscopy imaging of gel-specimen composites is still arduous. This research thus proposes an attainable Ex-dSTORM workflow through sample preparation, in-situ drift correction, optimized two-color imaging, and ultrastructural image analysis. Together, our Ex-dSTORM analyses provide a pragmatic roadmap to successfully determine the DAP organization and uncover the DAP root architecture in mammalian cells at ~3-nm precision, offering an unprecedentedly ultra-detailed framework of DAPs.

## Methods

### Reagents

Sodium acrylate (SA, 97%, 408220, Sigma-Aldrich), Acrylamide (AA, 40%, A4058, Sigma-Aldrich), Acrylamide/Bis-acrylamide (30%, 29:1, 1610156, Bio-Rad), N,N,N’,N’-Tetramethylethylenediamine (TEMED, 1610801, Bio-Rad), Ammonium persulfate (APS, 1610700, Bio-Rad), Paraformaldehyde (FA, 16%, 15710, Electron Microscopy Sciences), Sodium dodecyl sulfate (SDS, 0227, VWR Life science), Sodium chloride (NaCl, 31434, Sigma-Aldrich), Tris (1.5M, pH 8.8, J831, VWR Life science), Bovine serum albumin (BSA, A9647, Sigma-Aldrich), Tween 20 (P137, Sigma-Aldrich), Methyl alcohol (methanol, 15306121, Macron), Phosphate buffered saline (10X PBS, 70011044, Gibco), Dimethyl sulfoxide (DMSO, D8418, Sigma-Aldrich)

### Cell culture

Human retinal pigment epithelial cells (hTERT RPE-1, ATCC-CRL-4000) were cultured in Dulbecco’s modified Eagle’s medium (DMEM)/F-12 mixture medium with L-glutamine and HEPES (1:1; 11330-032, Gibco, Thermo Fisher Scientific) at 37°C under 5% CO_2_ with 10% fetal bovine serum (FBS, SH3010903, Hyclone), sodium bicarbonate (NaHCO3, S6014, Sigma-Aldrich), and 1% penicillinstreptomycin. Prior to fixation, RPE-1 cells were cultured on poly-L-lysine coated coverslips and cilium formation was induced by 24-48 h serum starvation.

### Antibodies

Detailed information on the primary antibodies used in this work is listed in Supplementary Table 5. Second antibodies used in this study were Alexa Fluor 647 (anti-mouse A31571, anti-rabbit A31573, anti-rat A21247; Thermo Fisher Scientific), Atto 647N (anti-mouse 50185-1ML-F; Sigma-Aldrich), CF568 (anti-rabbit 20098, anti-rat 20092; Biotium), and Alexa Fluor 488 (anti-mouse A21202; Sigma-Aldrich). Dyomics 654 (Dy654)-conjugated secondary antibodies were custom-made by conjugating Dy654 N-hydroxysuccinimidyl (NHS) ester (654-01; Dyomics) to different IgG antibodies respectively (anti-mouse 715-005-151, anti-rabbit 711-005-152, anti-rat 712-005-153; Jackson ImmunoResearch).

### CRISPR construction of CEP128^-/-^ cells

In order to inactivate CEP128 in RPE-1 cells, as described previously^47^, the RNA-guided targeting of genes was achieved through coexpression of the Cas9 protein with gRNAs using reagents from the Church group^48^ (Addgene: http://www.addgene.org/crispr/church/). Targeting sequences of the gRNAs were as followed: CEP128 gRNA2 (5’-GCTGCCAGATCAACGCACAGGG-3’), CEP128 gRNA4 (5’-GAGTCAGCTCTGAGATCTGAAGG-3’), CEP128 gRNA5 (5’-GCAGCTGAACTTCAGCGCAATGG-3’). To achieve complete protein depletion, we adopted multiple gRNAs targeting different exons. These targeting sequences were cloned into the gRNA vector (plasmid 41824, Addgene) via the Gibson assembly method (New England Biolabs). Pure CEP128 knockout cells were obtained through clonal propagation from a single cell and then confirmed by genotyping and immunoblotting as in our previous research^23^.

### Ultrastructure expansion microscopy protocol

RPE-1 cells on 12 mm coverslips were fixed with methanol at −20 °C for 10 min. Cells were then incubated in the perfusion solution containing 1.4% FA and 2% AA in PBS for 5-6 h at 37 °C. Next, gelation was proceeded via incubating each coverslip with U-ExM monomer solution (19% (w/w) SA, 10% (w/w) AA, 0.1% (w/w) BIS in PBS) supplemented with 0.5% TEMED and 0.5% APS in order in a pre-cooled humidified chamber. After prepolymerization on the ice for 1 min, the chamber was transferred into the incubator for gelation at 37 °C for one hour. Coverslips with hydrogel were placed in fresh denaturation buffer (200mM SDS, 200mM NaCl in 50mM Tris (pH8.8)) for 15 min at RT. Detached hydrogels were incubated in fresh denaturation buffer at 95 °C for 1.5 h.

After denaturation, the hydrogels were expanded in fresh double deionized water (ddH_2_O) at least three times (30 min for each) with gentle shaking and then incubated overnight in ddH_2_O. Next, the hydrogels were kept in PBS before immunostaining. Immunostaining was first carried out via staining the hydrogels with primary antibodies diluted in 2% BSA/PBS at 37 °C for three hours with gentle shaking, followed by washing three times with 0.1% Tween 20 in PBS and twice with PBS for 20 min each. Next, the hydrogels were stained with secondary antibodies diluted in 2% BSA/PBS at 37 °C for three hours with gentle shaking followed by the same washing steps. Finally, the hydrogels were expanded in the ddH_2_O until reaching their maximal expansion via exchanging ddH_2_O at least three times.

### Re-embedding of expanded hydrogels

A neutral acrylamide gel was crosslinked across the expanded hydrogel for stabilization in the dSTORM imaging buffer. The expanded hydrogels were incubated in freshly prepared re-embedding gel solution (10% (w/w) AA, 0.15% (w/w) BIS, 0.05% (w/w) TEMED, 0.05% (w/w) APS in ddH_2_O) twice for 25 min each time at RT with gentle shaking. The polymerization process was facilitated by placing the whole setup in a nitrogen-filled humidified chamber and incubating at 37 °C for 1.5 h. After polymerization, the re-embedding gels were washed three times in ddH_2_O for 20 min each.

### Determination of expansion factor

SCLT1 was utilized as the reference to determine the expansion factor after re-embedding through this research to determine the microscopic expansion factor. Nearly perfect top-view of SCLT1 were imaged via dSTROM and Ex-dSTORM, respectively. The expansion factor is then determined by the ratio of the mean diameter of SCLT1 between Ex-dSTORM and dSTORM.

### Ex-dSTORM imaging

Ex-dSTORM image acquisition was performed on a custom-built setup based on a commercial inverted microscope (Eclipse Ti-E, Nikon) with a focus stabilizing system and a laser merge module (ILE, Spectral Applied Research) with individual controllers for four light sources. For wide-field illumination of samples, beams from a 637 nm laser (OBIS 637 LX 140 mW, Coherent), a 561 nm laser (Jive 561 150 mW, Cobolt), a 488 nm laser (OPSL 488 LX 150 mW, Coherent), and a 405 nm laser (OBIS 405 LX 100 mW, Coherent) were homogenized (Borealis Conditioning Unit, Spectral Applied Research) and focused onto the back focal plane of an oil-immersing objective (100X 1.49, CFI Apo TIRF, Nikon). During Ex-dSTORM image acquisition, the 637 nm and 561 nm laser lines were operated at a high intensity of ~3 kW/cm^2^ to quench most of the fluorescence from Alexa Fluor 647, Dy654, and CF568. A weak 405 nm laser beam was introduced to convert a portion of fluorophores from a dark to a fluorescent state. The 488 nm laser line was intermittently switched on every 800 frames for in-situ drift correction. The emitted fluorescent signal was filtered through a quad-band filter (ZET405/488/561/640 mv2, Chroma), and then the signal was recorded on an electron-multiplying charge-coupled device (EMCCD) camera (Evolve 512 Delta, Photometrics) with a pixel size of 93 nm. For single-color imaging, signals of Alexa Fluor 647 were filtered with a quad-band filter; for two-color imaging, the Dy654 channel was first obtained, then the CF568 channel was acquired with a combination of quad-band and short-pass filter (BSP01-633R-25, Semrock). Generally, for each dSTORM image, 15,000-30,000 frames were acquired at a rate of 50 fps. The position of the individual single-molecule peak was then localized using MetaMorph Superresolution Module (Molecular Devices) based on a wavelet segmentation algorithm. In the figures, superresolution images were cleaned with the gaussian filter of 0.7-1 pixel.

Immobilized re-embedding hydrogels within a customized holder were immersed in an imaging buffer containing Tris-HCl, NaCl (TN) buffer at pH 8.0, 10-100 mM mercaptoethylamine (MEA) at pH 8.0, and an oxygen-scavenging system consisting of 10% glucose (G5767, Sigma-Aldrich), 0.5 mg mL^-1^ glucose oxidase, and 40 μg mL^-1^ catalase.

### In-situ drift correction

To perform in-situ drift correction, marker protein (ATP synthase in this study, ab109867, Abcam) was first labeled with Alexa Fluor 488 at a dilution of 1/100 before imaging. Lateral position drift was intermittently measured during acquisition and corrected by ImageJ. The sets of images with maker signals were eliminated before localizing each single-molecule peak. Chromatic aberration between red and far-red channels was compensated with a customized algorithm relocating each pixel of a 561 nm image to its targeted position in the 647 nm channel with a predefined correction function obtained by a parabolic mapping of multiple calibration beads.

### Imaging analysis

For axial imaging, a ring pattern of proteins in this research was used to identify the top view orientation. To determine the diameter of these proteins with minimal variations caused from batches to batches, cells to cells, or blades to blades of the sample, we used SCLT1 as the reference to measure the radius ratio of proteins (**Supplementary Fig. 9**) and scaled accordingly. For dispersed radial distribution of CEP164, radial positions were described with the radius ratio of each punctum with respect to SCLT1. To determine the relative angular position among DAP proteins, we estimated angle differences of puncta between the proteins and

SCLT1. More details are illustrated in Supplementary **Fig. 6**. For lateral-view imaging, a rod-like pattern was used to ensure the orientation. To determine the relative longitudinal position among DAP proteins, we quantified relative longitudinal distances between each punctum of proteins and SCLT1 (**Supplementary Fig. 9**). All the images of expanded samples were scaled by the expansion factor of 3.92. The angular correlation in **Fig. 6b** was determined based on the maximum correlation coefficient between C2CD3 and Ac-Tub signals. The same procedure was performed in **Fig. 7g** of the ODF2-N/SCLT1 protein pair due to its wider distribution in the circumferential direction.

### Three-dimensional model illustration

The 3D model of the DAP was constructed with the 3D illustration software Blender (Blender Foundation). The dimensions of the DAP model were based on the mean protein positions obtained from Ex-dSTORM analyses in the results (Supplementary Table 2-4). The ultra-detailed information of DAP molecular localizations from each protein was then applied to the DAP model correspondingly. The 3D model of the centriolar microtubules was based on the information from the previous study^49^.

## Acknowledgments

We thank Jung-Chi Liao and Meng-Fu Bryan Tsou for sharing reagents and cell lines. This work was supported by the Ministry of Science and Technology, Taiwan (Grant No. 109-2222-E002-003-MY3 and Grant No. 110-2628-E-002-012-) to T.T.Y.

## Author contributions

T.T.Y. and T.-J.B.C. conceived the study. T.-J.B.C. and J.C.H performed the Ex-dSTORM imaging and analyzed the data. T.T.Y. provided guidance and assistance on the project. T.-J.B.C. and T.T.Y. created the 3D computational model. T.-J.B.C., J.C.H., and T.T.Y. interpreted the data and wrote the manuscript.

## Data availability

All the data supporting the findings described in this study are available within the article and its supplementary information file or from the corresponding author upon reasonable request.

## Competing interests

The authors declare no competing interests.

## Notes

### Competing Interest Statement

The authors have declared no competing interest.

## References

1. Vorobjev I, YuS C. Centrioles in the cell cycle. I. Epithelial cells. The Journal of cell biology 93, 938–949 (1982).

2. Sorokin S. Reconstructions of centriole formation and ciliogenesis in mammalian lungs. Journal of cell science 3, 207–230 (1968).

3. Satir P, Christensen ST. Overview of structure and function of mammalian cilia. Annu Rev Physiol 69, 377–400 (2007).

4. Nigg EA, Raff JW. Centrioles, centrosomes, and cilia in health and disease. Cell 139, 663–678 (2009).

5. Ishikawa H, Marshall WF. Ciliogenesis: building the cell’s antenna. Nature reviews Molecular cell biology 12, 222–234 (2011).

6. Eggenschwiler JT, Anderson KV. Cilia and developmental signaling. Annu Rev Cell Dev Biol 23, 345–373 (2007).

7. Joo K, et al. CCDC41 is required for ciliary vesicle docking to the mother centriole. Proceedings of the National Academy of Sciences 110, 5987–5992 (2013).

8. Kobayashi T, Dynlacht BD. Regulating the transition from centriole to basal body. Journal of Cell Biology 193, 435–444 (2011).

9. Schmidt KN, Kuhns S, Neuner A, Hub B, Zentgraf H, Pereira G. Cep164 mediates vesicular docking to the mother centriole during early steps of ciliogenesis. Journal of Cell Biology 199, 1083–1101 (2012).

10. Tanos BE, et al. Centriole distal appendages promote membrane docking, leading to cilia initiation. Genes & development 27, 163–168 (2013).

11. Garcia-Gonzalo FR, Reiter JF. Scoring a backstage pass: mechanisms of ciliogenesis and ciliary access. Journal of Cell Biology 197, 697–709 (2012).

12. Reiter JF, Blacque OE, Leroux MR. The base of the cilium: roles for transition fibres and the transition zone in ciliary formation, maintenance and compartmentalization. EMBO reports 13, 608–618 (2012).

13. Rosenbaum JL, Witman GB. Intraflagellar transport. Nature reviews Molecular cell biology 3, 813–825 (2002).

14. Bornens M. Centrosome composition and microtubule anchoring mechanisms. Current opinion in cell biology 14, 25–34 (2002).

15. Graser S, et al. Cep164, a novel centriole appendage protein required for primary cilium formation. The Journal of cell biology 179, 321–330 (2007).

16. Sillibourne JE, Hurbain I, Grand-Perret T, Goud B, Tran P, Bornens M. Primary ciliogenesis requires the distal appendage component Cep123. Biology open 2, 535–545 (2013)

17. Wei Q, et al. Transition fibre protein FBF1 is required for the ciliary entry of assembled intraflagellar transport complexes. Nature communications 4, 1–10 (2013).

18. Ye X, Zeng H, Ning G, Reiter JF, Liu A. C2cd3 is critical for centriolar distal appendage assembly and ciliary vesicle docking in mammals. Proceedings of the National Academy of Sciences 111, 2164–2169 (2014).

19. Lo C-H, et al. Phosphorylation of CEP83 by TTBK2 is necessary for cilia initiation. Journal of Cell Biology 218, 3489–3505 (2019).

20. Hoover AN, Wynkoop A, Zeng H, Jia J, Niswander LA, Liu A. C2cd3 is required for cilia formation and Hedgehog signaling in mouse. (2008).

21. Ishikawa H, Kubo A, Tsukita S, Tsukita S. Odf2-deficient mother centrioles lack distal/subdistal appendages and the ability to generate primary cilia. Nature cell biology 7, 517–524 (2005).

22. Tateishi K, et al. Two appendages homologous between basal bodies and centrioles are formed using distinct Odf2 domains. Journal of Cell Biology 203, 417–425 (2013).

23. Chong WM, et al. Super-resolution microscopy reveals coupling between mammalian centriole subdistal appendages and distal appendages. Elife 9, e53580 (2020).

24. Sillibourne JE, et al. Assessing the localization of centrosomal proteins by PALM/STORM nanoscopy. Cytoskeleton 68, 619–627 (2011).

25. Lau L, Lee YL, Sahl SJ, Stearns T, Moerner W. STED microscopy with optimized labeling density reveals 9-fold arrangement of a centriole protein. Biophysical journal 102, 2926–2935 (2012).

26. Yang TT, et al. Super-resolution architecture of mammalian centriole distal appendages reveals distinct blade and matrix functional components. Nature communications 9, 1–11 (2018).

27. Bowler M, et al. High-resolution characterization of centriole distal appendage morphology and dynamics by correlative STORM and electron microscopy. Nature communications 10, 1–15 (2019).

28. Tsai J-J, Hsu W-B, Liu J-H, Chang C-W, Tang TK. CEP120 interacts with C2CD3 and Talpid3 and is required for centriole appendage assembly and ciliogenesis. Scientific reports 9, 1–15 (2019).

29. Wang L, Failler M, Fu W, Dynlacht BD. A distal centriolar protein network controls organelle maturation and asymmetry. Nature communications 9, 1–15 (2018).

30. Chen F, Tillberg PW, Boyden ES. Expansion microscopy. Science 347, 543–548 (2015).

31. Tillberg PW, et al. Protein-retention expansion microscopy of cells and tissues labeled using standard fluorescent proteins and antibodies. Nature biotechnology 34, 987–992 (2016).

32. Chozinski TJ, et al. Expansion microscopy with conventional antibodies and fluorescent proteins. Nature methods 13, 485–488 (2016).

33. Ku T, et al. Multiplexed and scalable super-resolution imaging of three-dimensional protein localization in size-adjustable tissues. Nature biotechnology 34, 973–981 (2016).

34. Chen F, et al. Nanoscale imaging of RNA with expansion microscopy. Nature methods 13, 679–684 (2016).

35. Alon S, et al. Expansion sequencing: Spatially precise in situ transcriptomics in intact biological systems. Science 371, eaax2656 (2021).

36. Gao R, et al. Cortical column and whole-brain imaging with molecular contrast and nanoscale resolution. Science 363, eaau8302 (2019).

37. Gambarotto D, et al. Imaging cellular ultrastructures using expansion microscopy (U-ExM). Nature methods 16, 71–74 (2019).

38. Halpern AR, Alas GC, Chozinski TJ, Paredez AR, Vaughan JC. Hybrid structured illumination expansion microscopy reveals microbial cytoskeleton organization. ACS nano 11, 12677–12686 (2017).

39. Gao M, et al. Expansion stimulated emission depletion microscopy (ExSTED). ACS nano 12, 4178–4185 (2018).

40. Xu H, et al. Molecular organization of mammalian meiotic chromosome axis revealed by expansion STORM microscopy. Proceedings of the National Academy of Sciences 116, 18423–18428 (2019).

41. Zwettler FU, et al. Molecular resolution imaging by post-labeling expansion single-molecule localization microscopy (Ex-SMLM). Nature communications 11, 1–11 (2020).

42. Thauvin-Robinet C, et al. The oral-facial-digital syndrome gene C2CD3 encodes a positive regulator of centriole elongation. Nature genetics 46, 905–911 (2014).

43. Viol L, et al. Nek2 kinase displaces distal appendages from the mother centriole prior to mitosis. Journal of Cell Biology 219, (2020).

44. Kuhns S, et al. The microtubule affinity regulating kinase MARK4 promotes axoneme extension during early ciliogenesis. Journal of Cell Biology 200, 505–522 (2013).

45. Zwettler FU, et al. Tracking down the molecular architecture of the synaptonemal complex by expansion microscopy. Nature communications 11, 1–11 (2020).

46. Helmerich DA, Beliu G, Matikonda SS, Schnermann MJ, Sauer M. Photoblueing of organic dyes can cause artifacts in super-resolution microscopy. Nature Methods 18, 253–257 (2021).

47. Mazo G, Soplop N, Wang W-J, Uryu K, Tsou M-FB. Spatial control of primary ciliogenesis by subdistal appendages alters sensation-associated properties of cilia. Developmental cell 39, 424–437 (2016).

48. Esvelt KM, Mali P, Braff JL, Moosburner M, Yaung SJ, Church GM. Orthogonal Cas9 proteins for RNA-guided gene regulation and editing. Nature methods 10, 1116–1121 (2013).

49. Anderson RG. The three-dimensional structure of the basal body from the rhesus monkey oviduct. The Journal of cell biology 54, 246–265 (1972).

