## Supporting information for "Single-molecule localization microscopy reveals the ultrastructural root constitution of distal appendages in expanded mammalian centrioles"

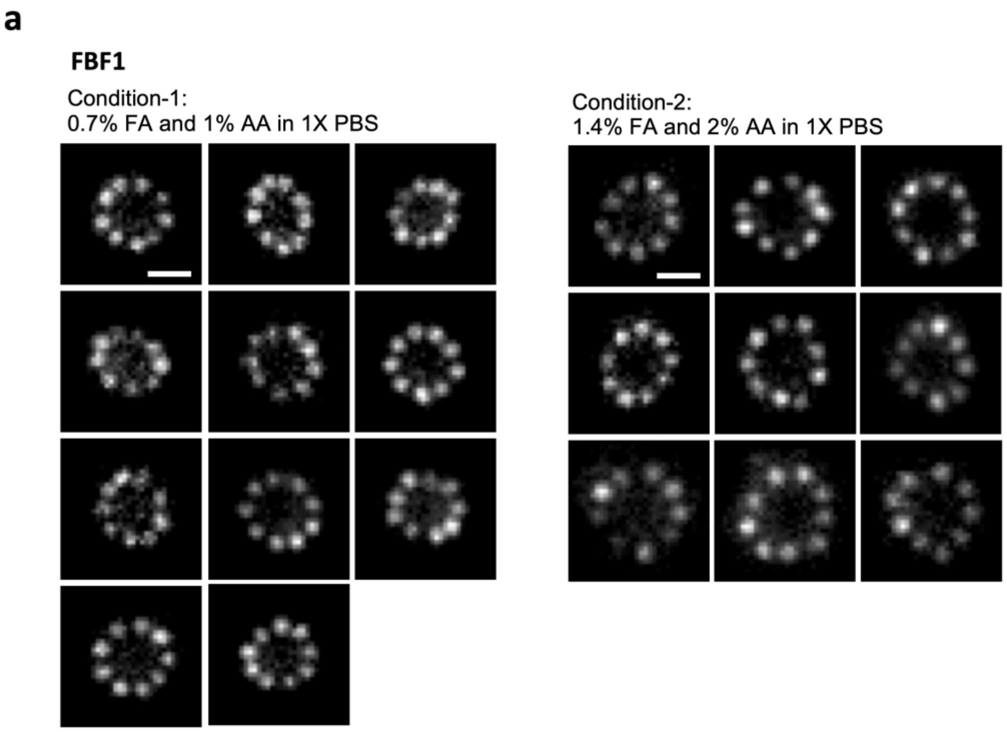


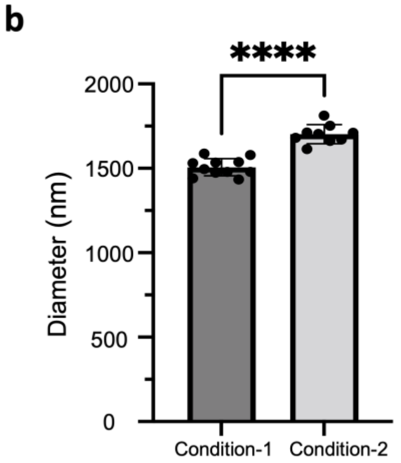


**c**

**Diameter of FBF1**

| | Mean diameter (nm) | Expansion factor |
|---|---|---|
| Condition-1 | 1506 ± 51 (n=11) | 3.51 |
| Condition-2 | 1702 ± 57 (n=9) | 3.97 |
| Rate of increase (volume) | 44 % | |

**Supplementary Figure 1. Optimizing concentration of perfusion step enhancing the expansion factor. a** ExM images of FBF1 under two perfusion conditions. Condition-2 doubles FA and AA concentration compared with the original recipe (condition-1), which facilitates a higher expansion factor. **b** Statistical analysis with the significant difference of mean diameter under two concentrations. ****$p<0.0001$, unpaired two-tailed t-test. **c** Table of mean diameter under two conditions with corresponding expansion factors compared with our previous result[1]. Scale bar, 1 μm (**a**).

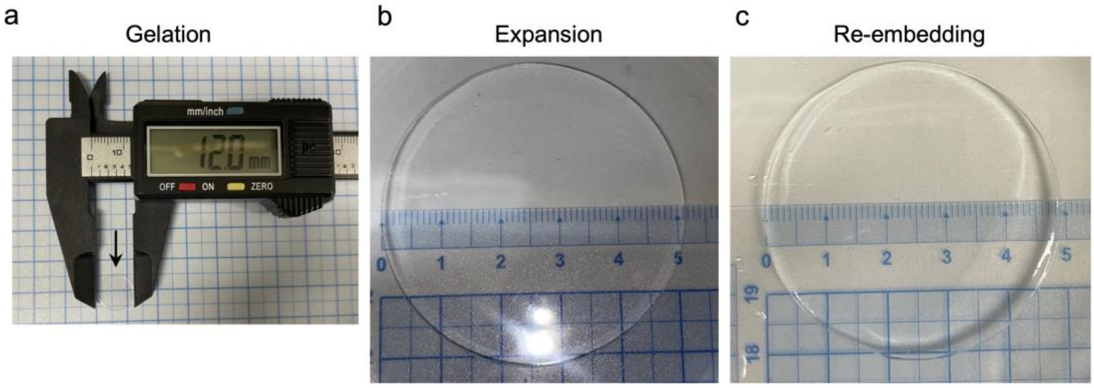


d

| | Gelation | Expansion | Re-embedding |
|---|---|---|---|
| Mean diameter (mm) | 12.0 ± 0.0 | 51.7 ± 0.6 | 48.2 ± 0.3 |
| Expansion factor | 1 | 4.3 | 4.0 |

**Supplementary Figure 2. Determination of the expansion factor. a** Measurement of the diameter of the sample after gelation. **b** Measurement of the diameter of the expanded hydrogel. **c** Measurement of the diameter of the hydrogel after re-embedding. **d** Table of mean diameter of the hydrogel in different conditions with the determination of expansion factor after re-embedding (three independent experiments).

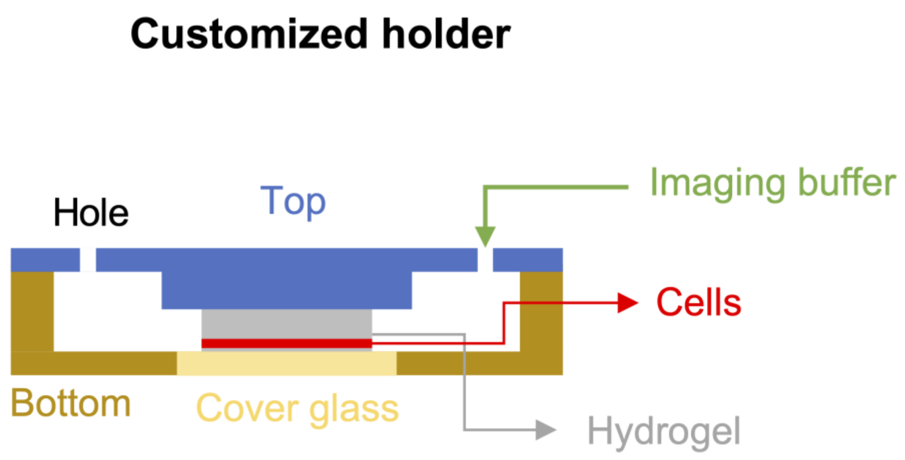


**Supplementary Figure 3. Schematic diagram of the customized sample holder for Ex-dSTORM imaging.** By sandwiching the hydrogel after re-embedding, we can minimize lateral drift without chemical modification of the cover glass. Two small holes on both sides of top panel are prepared for filling imaging buffer and preventing the interaction between the imaging buffer and the external environment.

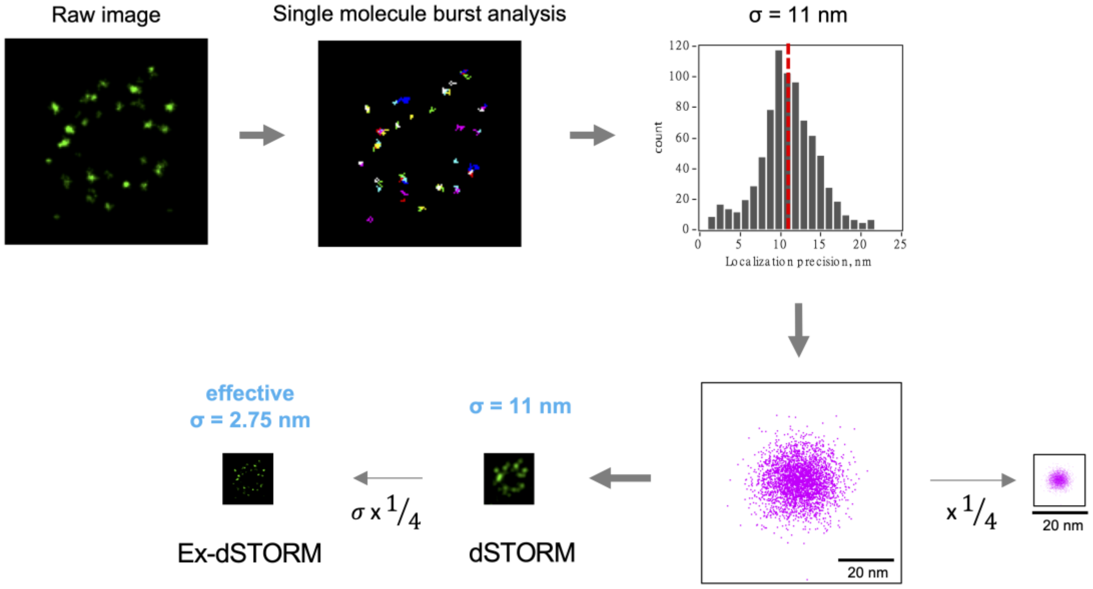


**Supplementary Figure 4. Validation of our Ex-dSTORM resolution.** For evaluation of resolution in Ex-dSTORM, we utilize an Ex-dSTORM image to evaluate the resolution of our system. Histogram analysis reveals the mean localization precision of the image is equal to 11 nm by measuring the localization precision per switching event. By dividing the localization precision with expansion factor in **Supplementary Fig. 1d**, the effective localization precision of Ex-dSTORM imaging is approximately to 3 nm.

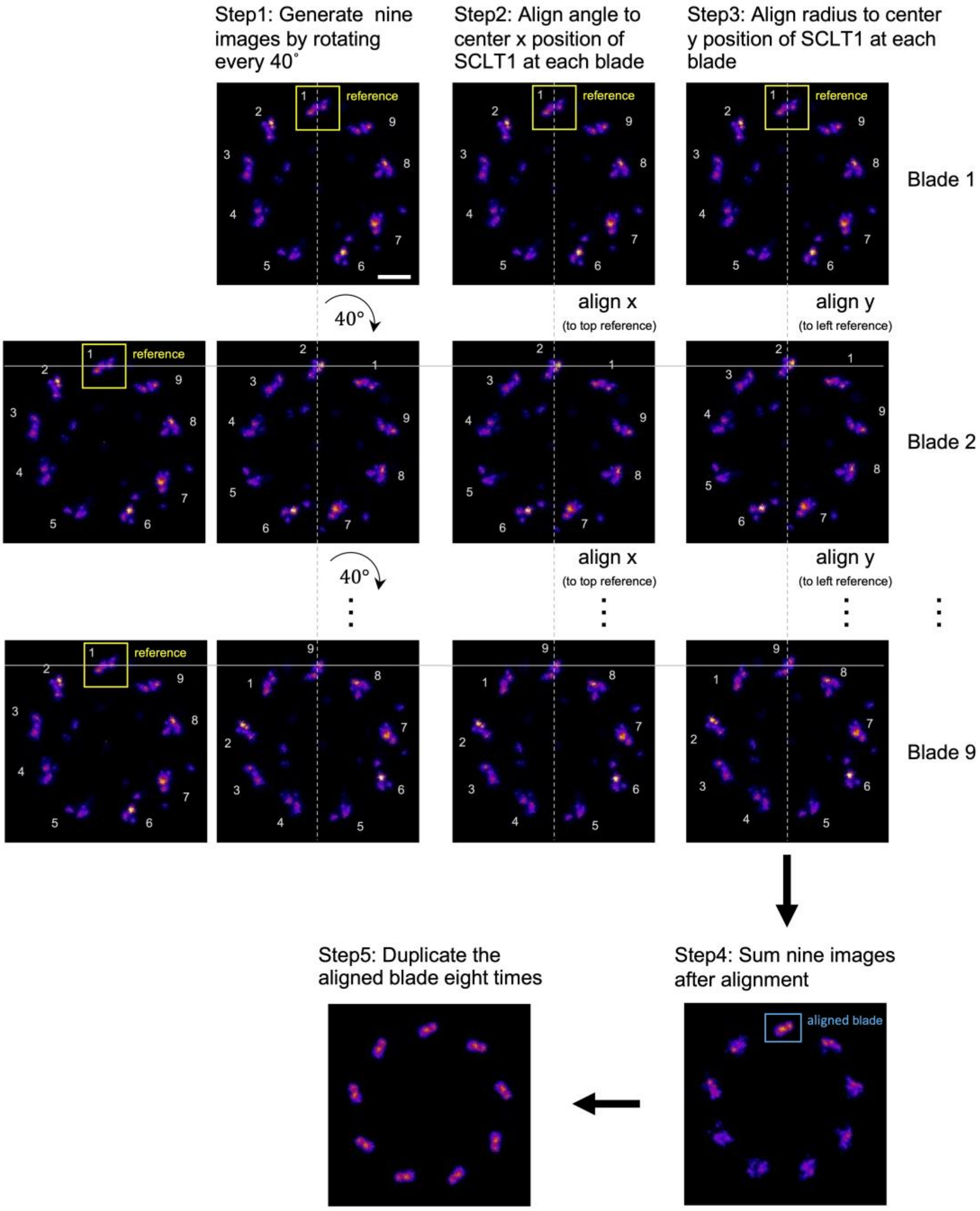


**Supplementary Figure 5. Pipeline of averaging Ex-dSTORM signals (refer to Fig. 1e).** To strengthen the ultra-detailed features of the proteins, we average the Ex-dSTORM signals with representative images. First, we rotate the original Ex-dSTORM image every 360˚/9 around its geometrical center (9 images). Second, we align the angle to the center x position of SCLT1 at each blade. Third, we perform the longitudinal movement of every image to align its center of blade compared with the reference blade. Fourth, we sum nine aligned images together to extract the features of individual protein (top aligned blade). Finally, we duplicate the aligned blade from step 4 eight times with a 40˚ interval to acquire the final averaged image with ultra-detailed features. Scale bar, 100 nm.

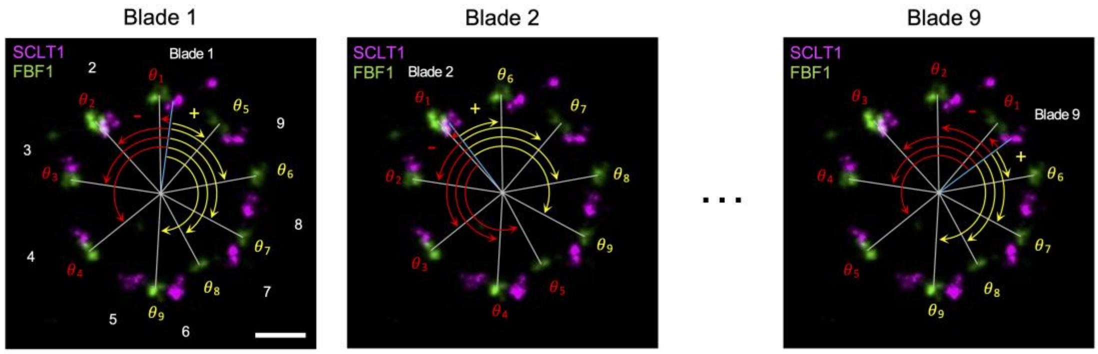


**Supplementary Figure 6. Measurement of relative angles.** We apply the angular analysis to determine relative radial direction among proteins (observed from the distal-view of the centriole). The angular positions of each protein punctum are first recorded in the Ex-dSTORM images. The difference in angles is then determined from the center of reference (blue line) to the center of targets (gray lines) and assigned with the direction (minus sing: counterclockwise, plus sign: clockwise). Angle differences for a single reference are obtained by calculating the included angles between the reference (blue line) and the targets (gray lines) from the nearest target signals ($\theta_1$) all the way to the farthest signals ($\theta_4$ and $\theta_9$). The rest of angle differences for other references in distinct blades are measured in the same manner (Blade1 to Blade 9). Scale bar, 100 nm.

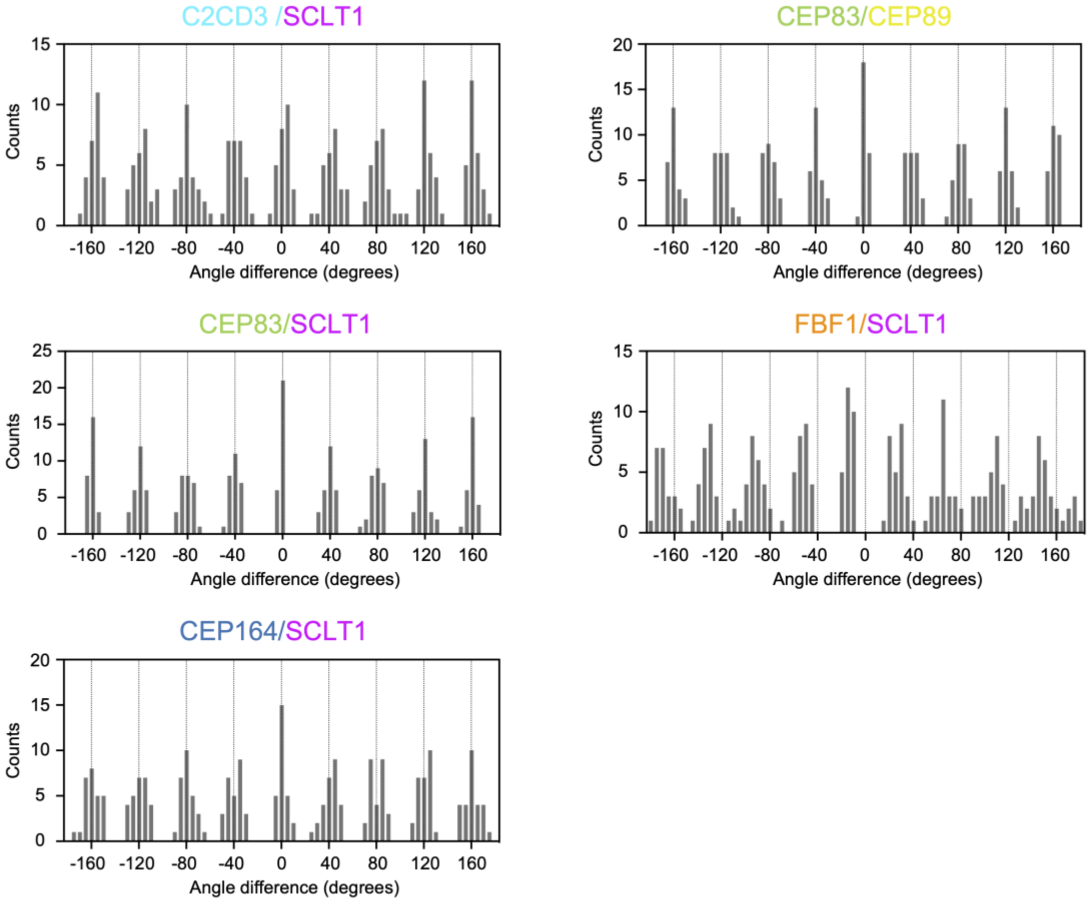


**Supplementary Figure 7. Angular analyses in the radial direction revealing the protein-protein angular relationship.** The histogram analysis of angle differences of various protein pairs reveals the angular spacing and the relative angle. In all protein pairs, the angular spacings between blades are found to be the multiples of 40˚. All pairs except FBF1-SCLT1 pair, the relative angle between blades are the multiples of 40˚, indicating these proteins are located in the same radial direction in a single blade. As for FBF1-SCLT1 pair, the result demonstrates that FBF1 is positioned in the distinctive radial direction against SCLT1 and also other DAP proteins.

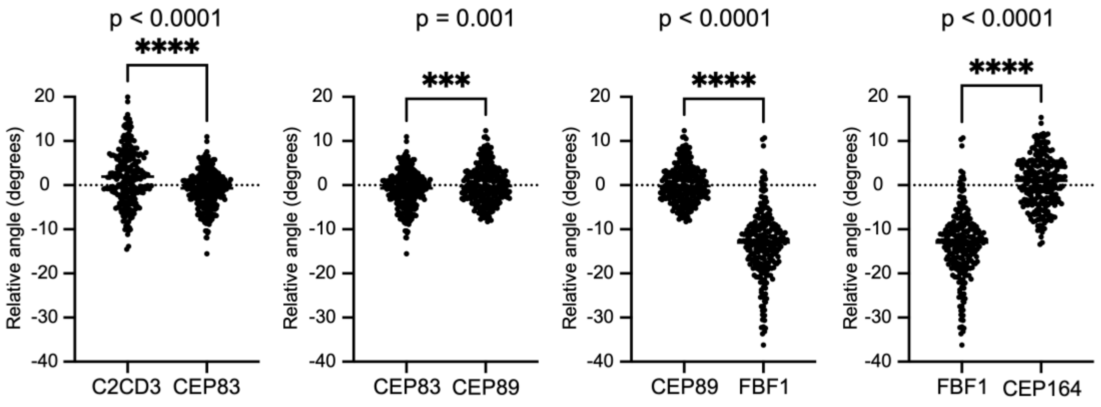


**Supplementary Figure 8. Statistical analysis of the relative angle among DAP proteins.** The relative angles between DAP proteins and SCLT1 are demonstrated in the above plots. The statistical analyses reveal the statistical significance of the relative angles among different DAP proteins (unpaired two-tailed t-test).

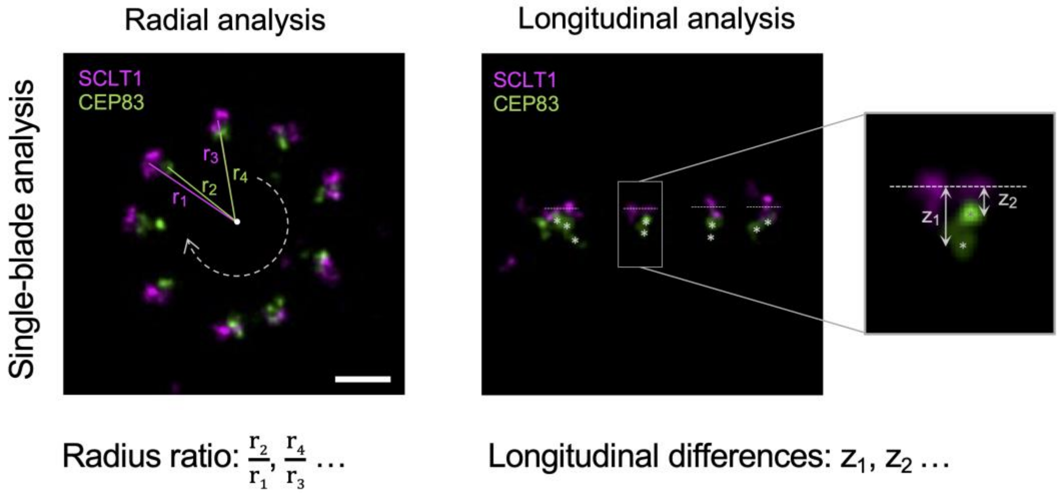


**Supplementary Figure 9. Single-blade resolution for radial and longitudinal analyses.** In radial analysis, we calculated the radius ratio but not the real values, eliminating the possible blade-to-blade and cell-to-cell variation resulted from different batches of the sample after re-embedding. The radius ratios are calculated in a single blade by dividing the center of target signals (CEP83) against the center of reference signals (SCLT1) and the same processes are repeated in all the blades. In longitudinal analysis, the longitudinal differences are obtained by subtracting the longitudinal position of the reference center (dashed lines) from that of every punctum of targets (asterisks) in a single blade ($z_1$, $z_2$, inset). To prevent the possible variations in the longitudinal analysis, we have checked the mean diameter of SCLT1 in every batch of the sample within 5% difference. Scale bar, 100 nm.

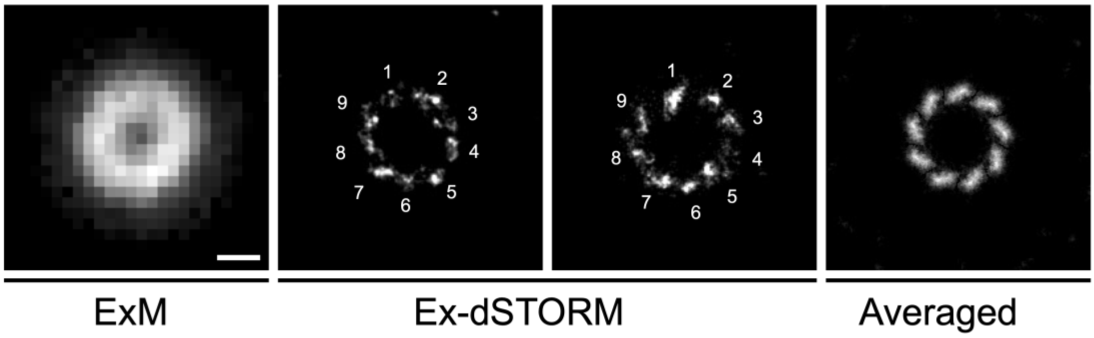


**Supplementary Figure 10. Ex-dSTORM resolving nine discrete microtubule triplets signals.** With the resolution under Ex-dSTORM, we could perspicuously resolve the characteristic nine microtubule triples. The averaged image of every 40˚ rotation enhances the features of microtubule triplets. Scale bar, 100 nm.

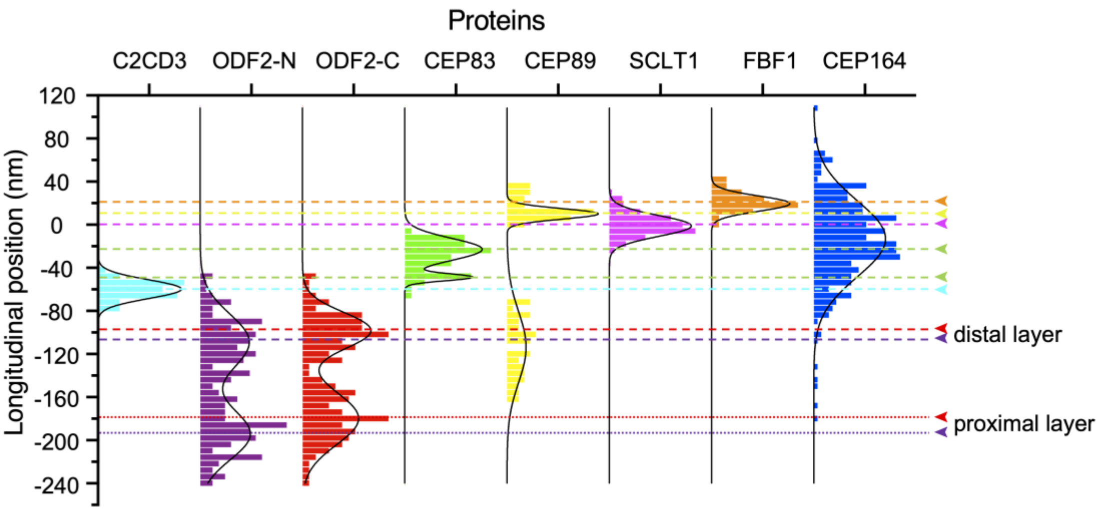


**Supplementary Figure 11. Histogram analysis of the longitudinal positions of DAP proteins and ODF2-C/N relative to SCLT1.** Mean longitudinal position of each protein are marked with arrowheads (n ≥ 7 centrioles for each). Two-layered distributions of ODF2-C/N can be clearly identified in the analysis. Distal-ODF2-C/N are found to be slightly lower than other core DAP proteins, indicating a pedestal of other DAP proteins outside the mother centriole.

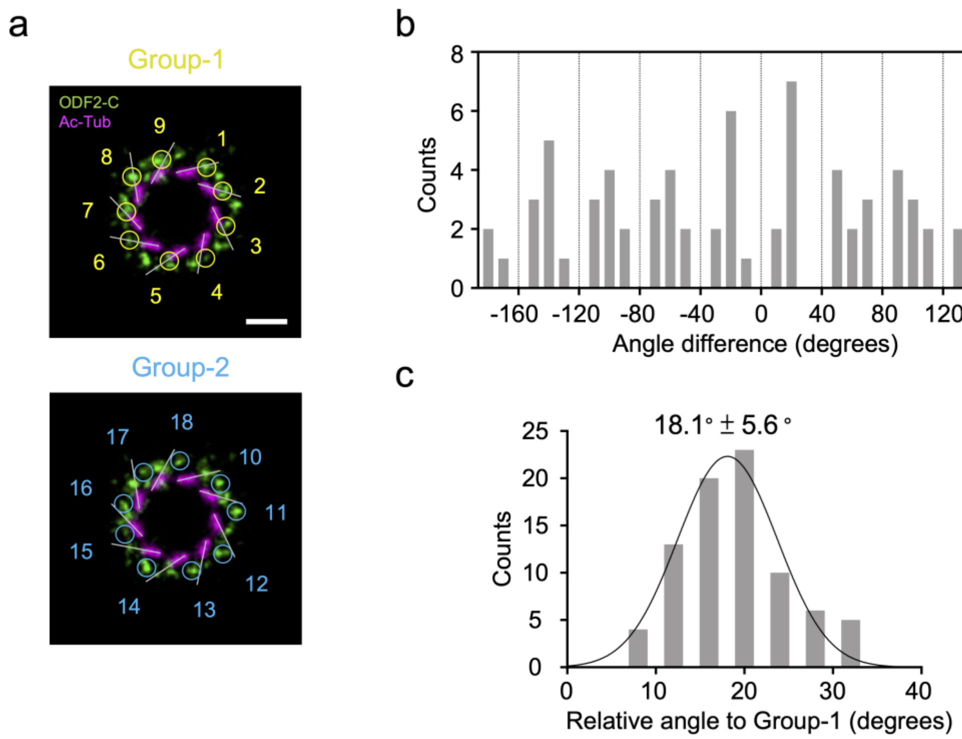


**Supplementary Figure 12. Quantification of the angular offset between two groups of ODF2. a** Representative image of two groups populations of ODF2 with Ac-Tub as the reference. **b** Histogram analysis of relative angle between both group-1 and group-2 of ODF2, utilizing group-1 of ODF2 as the reference and the analytical method in **Supplementary Fig. 6** and **7**. **c** Gaussian fitting with all measured points revealing an 18° clockwise offset of group-2 of ODF2 relative to group-1 of ODF2.

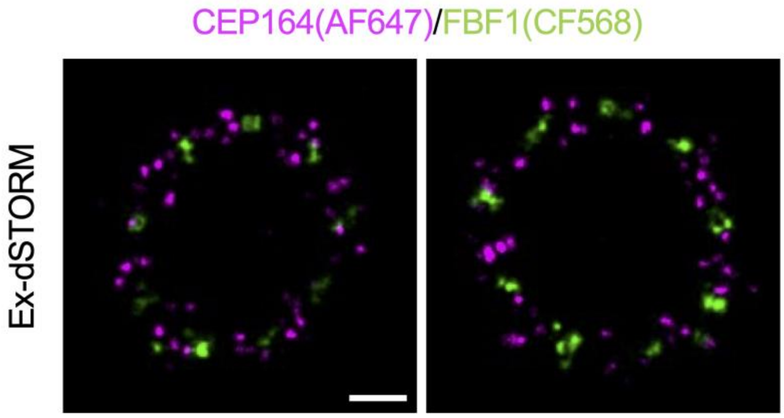


**Supplementary Figure 13. Representative Ex-dSTORM images showing negligible crosstalk using AF647.** Images of protein pair of CEP164 (AF647) and FBF1 (CF568) with distinct angular distributions along circumferential direction demonstrates acceptable crosstalk ratio when stain AF647 for the protein with lower protein numbers. Scale bar, 100nm.

**Supplementary Table 1 Determination of the expansion factor via dividing the mean diameter of SCLT1 analyzed from Ex-dSTORM images by that from dSTORM images**

| SCLT1 | Diameter (mean ± sd, nm) | Expansion factor |
|---|---|---|
| dSTORM | 402.0 ± 20.7 (108 BLs) | – |
| Ex-dSTORM | 1576.1 ± 128.0 (135 BLs) | 1576.1/402.0 = 3.92 |

(BL = blade)

**Supplementary Table 2 Ultra-detailed analyses of core DAP proteins**

| Proteins | Characteristic length (XY) (mean ± sd, nm) | Rotation angle (mean ± sd, degrees) | Longitudinal distance (Z) (mean ± sd, nm) | Inclination angle (mean ± sd, degrees) * |
|---|---|---|---|---|
| C2CD3 | – | broad distribution (31BLs) | – | – |
| CEP83 | 20.2 ± 3.3 (58 BLs) | 30.4 ± 11.6° (45 BLs) | 25.9 ± 5.5 (36 BLs) | 52.1 ± 19.0° |
| CEP89 | 29.3 ± 5.5 (32 BLs) | 51.3 ± 10.9° (38 BLs) | 11.2 ± 5.0 (15 BLs) | 21.0 ± 9.7° |
| SCLT1 | 31.4 ± 5.5 (55 BLs) | 55.5 ± 13.0° (41 BLs) | 8.6 ± 3.9 (25 BLs) | 16.4 ± 8.1° |
| FBF1 | 30.0 ± 5.5 (49 BLs) | broad distribution (43 BLs) | 10.6 ± 4.6 (20 BLs) | 19.5 ± 10.1° |

(BL = blade)

*Inclination angles were calculated as the ratio of characteristic length to longitudinal distance.

**Supplementary Table 3 Ultra-resolved diameter, relative angle, and chirality of various proteins**

| Proteins | Diameter (mean ± sd, nm) | Relative angle (mean ± sd, degrees) | Chirality |
|---|---|---|---|
| C2CD3 | 143.9 ± 15.3 (36 BLs) | 1.7 ± 6.3° (3 MCs) | – |
| distal ODF2-N | 273.4 ± 20.1 (45 BLs) | 15 ± 4.2° (3 MCs) | – |
| distal ODF2-C | 307.1 ± 19.0 (36 BLs) | 1.8 ± 5.5° (3 MCs) | counterclockwise** |
| CEP83 | 357.8 ± 14.6 (36 BLs) | -0.6 ± 4.0° (3 MCs) | clockwise |
| CEP89 | 366.2 ± 19.5 (36 BLs) | 0.2 ± 4.5° (3 MCs) | clockwise |
| SCLT1 | 402.2 ± 32.6 (126 BLs) | 0° | clockwise |
| FBF1 | 408.4 ± 25.7 (36 BLs) | -12.9 ± 5.3° (3 MCs) | – |
| CEP164 | 417.7 ± 76.3 (36 BLs)* | 0.1 ± 6.2° (3 MCs) | counterclockwise |
| Ac-Tub | 200.7 ± 14.7 (45 BLs) | 22.2 ± 7.0° (6 MCs) | clockwise |

(BL = blade, MC= mother centriole)

*Radial distribution of CEP164: 265.1 – 570.3 nm (broad distribution denoted as mean ± 2 sd)

**44% of ODF2-C represents counterclockwise chirality (48/108 BLs)

**Supplementary Table 4 Ultra-resolved longitudinal locations of various proteins**

| Proteins | Longitudinal position relative to SCLT1 (mean ± sd, nm) | Number of measured points |
|---|---|---|
| C2CD3 | -60.0 ± 8.7 | 52 (9 MCs) |
| ODF2-N | -109.1 ± 26.0; -194.5 ± 23.6 | 205 (6 MCs) |
| ODF2-C | -98.3 ± 16.8; -180.0 ± 24.4 | 161(6 MCs) |
| CEP83 | -23.0 ± 10.3; -49.0 ± 3.2 | 64 (9 MCs) |
| CEP89 | 9.9 ± 4.9; -114 ± 33.3 | 86 (14 MCs) |
| SCLT1 | -1.3 ± 9.6 | 266 (24 MCs) |
| FBF1 | 19.4 ± 7.0 | 68 (13 MCs) |
| CEP164 | -15.9 ± 40.0* | 289 (9 MCs) |

(MC = mother centriole)

*Longitudinal distribution of CEP164: -95.9 – 64.1nm (broad distribution denoted as mean ± 2 sd)

### Supplementary Table 5 List of primary antibodies in the study

| Designation of antibody | Host | Source or reference | Immunogen (Sequence) | Identifiers | Dilution (Ex-dSTORM) |
|---|---|---|---|---|---|
| C2CD3 | rabbit IgG | Sigma-Aldrich | human C2CD3 (803-905 aa) | HPA040433 | 1/250 |
| Acetyl-alpha Tubulin (Ac-Tub) | mouse IgG | Thermo Fisher | – | 32-2700 | 1/200 |
| CEP83 | rabbit IgG | Sigma-Aldrich | human CEP83 (578-677 aa) | HPA038161 | 1/100 |
| CEP89 | rat IgG | Tanos et al. Genes & development (2013)[2] | – | – | 1/100 |
| SCLT1 | rat IgG | Tanos et al. Genes & development (2013)[2] | – | – | 1/100 |
| FBF1 | rabbit IgG | Proteintech | human FBF1 (20-347 aa) | 11531-1-AP | 1/100 |
| CEP164 | rabbit IgG | Proteintech | human CEP164 (1-112 aa) | 22227-1-AP | 1/250 |
| ODF2-C | rabbit IgG | Abcam | – | ab43840 | 1/100 |
| ODF2-N | rabbit IgG | Sigma-Aldrich | human ODF2 (39-200 aa) | HPA001874 | 1/100 |
| ATP synthase | mouse IgG | Abcam | – | ab109867 | 1/125 |
| CEP164 | goat IgG | Santa Cruz Biotechnology | human CEP164 N-terminus | sc-240226 | 1/100 |
| Polyglutamylated tubulin (GT335) | mouse IgG | AdipoGen | – | AG-20B0020-C100 | 1/200 |

(aa = amino acid)

### Supplementary references


1. Yang TT, *et al.* Super-resolution architecture of mammalian centriole distal appendages reveals distinct blade and matrix functional components. *Nature communications* **9**, 1-11 (2018).

2. Tanos BE, *et al.* Centriole distal appendages promote membrane docking, leading to cilia initiation. *Genes & development* **27**, 163-168 (2013).
